# Torsin ATPases influence chromatin interaction of the Torsin regulator LAP1

**DOI:** 10.1101/2020.10.01.321869

**Authors:** Naemi Luithle, Jelmi uit de Bos, Ruud Hovius, Daria Maslennikova, Renard Lewis, Rosemarie Ungricht, Beat Fierz, Ulrike Kutay

**Affiliations:** Institute of Biochemistry, Department of Biology, ETH Zurich, CH-8093 Zurich, Switzerland; Molecular Life Sciences Ph.D. Program, 8057 Zurich, Switzerland; Institute of Chemical Sciences and Engineering - ISIC, EPFL Lausanne, Switzerland; Celgene, Zürich, Switzerland; Novartis Institute of Biomedical Research, Basel, Switzerland

**Keywords:** nuclear envelope, inner nuclear membrane, mitosis, Torsins, chromosome segregation, cell division

## Abstract

The inner nuclear membrane is functionalized by diverse transmembrane proteins that associate with nuclear lamins and/or chromatin. When cells enter mitosis, membrane-chromatin contacts must be broken to allow for proper chromosome segregation; yet how this occurs remains ill-understood. Unexpectedly, we observed that an imbalance in the levels of the lamina-associated polypeptide 1 (LAP1), an activator of ER-resident Torsin AAA+-ATPases, causes a failure in membrane removal from mitotic chromatin, accompanied by chromosome segregation errors and changes in post-mitotic nuclear morphology. These defects are dependent on a hitherto unknown chromatin-binding region of LAP1 that we have delineated. LAP1-induced NE abnormalities are efficiently suppressed by expression of wild-type but not ATPase-deficient Torsins. Furthermore, a dominant-negative Torsin induces chromosome segregation defects in a LAP1-dependent manner. These results indicate that association of LAP1 with chromatin in the nucleus can be modulated by Torsins in the perinuclear space, shedding new light on the LAP1-Torsin interplay.

## Introduction

In all eukaryotes, a double membrane barrier termed nuclear envelope (NE) serves as the boundary of the nuclear compartment that safeguards the genetic information. The NE is built by a large chromatin-attached membrane sheet of the endoplasmic reticulum (ER). Despite the connectivity of the NE-ER membrane network, the inner nuclear membrane (INM) contains a unique set of transmembrane proteins (Holmer & Worman, 2001, Ungricht & Kutay, 2015). In general, enrichment of these membrane proteins from the peripheral ER at the nuclear face of the NE relies on several domains or short linear motifs in their extralumenal domains that together ensure retention on nuclear partners such as nuclear lamins, chromatin-associated factors or DNA (Boni et al., 2015, Powell & Burke, 1990, Ungricht et al., 2015). While the interaction of INM proteins with the nuclear lamina is restricted to lamin-expressing metazoan cells and certain protists, chromatin is the principal binding partner of INM proteins in all eukaryotes. The association of INM proteins with chromatin is functionally important, being it for the formation of gametes, for development or differentiation, exemplified by the NE-based pairing of homologous chromosomes during meiosis, or the progressive enrichment of transcriptionally repressed chromatin domains at the nuclear periphery of differentiating metazoan cells (Ungricht & Kutay, 2017).

How INM proteins associate with chromatin is best understood for cases in which folded domains are used for chromatin binding. The lamin B receptor (LBR), for example, contains an N-terminal tudor domain for interaction with the epigenetically marked tail of histone H4 as well as structurally less-defined regions for association with DNA and HP1 (Hirano et al., 2012). Some INM proteins contain short bihelical motifs for chromatin interaction such as the LEM (<u>L</u>AP2-<u>e</u>merin-<u>M</u>AN1) domain, which binds the chromatin factor BAF, or related motifs that bind DNA directly (Barton et al., 2015, Brachner & Foisner, 2011). The interaction of other mammalian INM proteins with chromatin is less understood. Biochemical experiments suggest that many INM proteins might even interact with DNA directly (Ulbert et al., 2006).

In cells undergoing open mitosis, the entire nuclear compartment is disintegrated by NE breakdown (NEBD), which liberates chromosomes from confinement by the nuclear membrane (Champion et al., 2017). In the course of prophase, the multifaceted interactions between INM proteins, chromatin and nuclear lamins are broken, allowing for the separation of membranes from chromatin and the partitioning of INM proteins into the mitotic ER network (Ellenberg et al., 1997, Yang et al., 1997). Exploiting a synthetic membrane–chromatin tethering system, we recently demonstrated that failed removal of membranes from chromatin impairs mitotic chromatin organization, chromosome segregation and cytokinesis, and perturbs post-mitotic nuclear morphology (Champion et al., 2019). Notably, also in yeasts with closed mitosis, the detachment of chromosomes from the NE is required for faithful chromosome segregation (Titos et al., 2014). It is generally assumed that changes in posttranslational modifications of INM proteins and chromatin-associated factors, especially protein phosphorylation, induce membrane dissociation from chromatin during mitotic entry. In strong support of this view, many mammalian INM proteins are phosphorylated in their nucleoplasmic domains by CDK1 or other abundant mitotic kinases during mitotic entry (Guttinger et al., 2009). Also chromatin-associated interaction partners of INM proteins such as BAF are released from chromatin by mitotic phosphorylation (Samwer et al., 2017). After prophase, multiple mechanisms ensure that the ER does not re-associate with chromatin prematurely, before NE reformation starts in late anaphase (Champion et al., 2019).

When studying the release of membrane proteins from chromatin during mitotic entry in cultured human somatic cells, we observed that an increased cellular level of one specific INM protein, the lamina-associated polypeptide 1 (LAP1), caused a failure in membrane removal from chromatin in mitosis. This observation not only indicated that LAP1 is a novel chromatin-binding INM protein but also suggested that some cellular factor(s) might have become limiting for breaking mitotic LAP1-chromatin contacts. LAP1 is an INM-localized activator of Torsins; the only known AAA+ ATPases residing in the lumen of the ER and the continuous perinuclear space (Brown et al., 2014, Goodchild & Dauer, 2005, Laudermilch & Schlieker, 2016, Sosa et al., 2014). Although neither the precise biological role nor the substrates of Torsins are known, loss of Torsin functionality causes changes in NE morphology (Goodchild et al., 2005), pointing towards a role at the NE. Mutations in *TOR1A*, the predominant Torsin isoform expressed in the brain (Jungwirth et al., 2010), lead to a severe movement disorder termed early-onset dystonia (DYT1) (Ozelius et al., 1997). Notably, also mutations in the *LAP1* gene, which is ubiquitously expressed in the human body, have been linked to primary dystonia as well as to cardiomyopathy, cerebellar atrophy and cancer (Dorboz et al., 2014, Fichtman et al., 2019, Kayman-Kurekci et al., 2014, Rebelo et al., 2015).

In our experiments, persistence of LAP1 on mitotic chromatin was accompanied by changes in post-mitotic NE morphology and cell division errors, reminiscent of defects that we had previously reported for a synthetic membrane–chromatin tether, which prevents mitotic chromatin release. Strikingly, overexpression of wild-type but not of ATPase-deficient Torsins efficiently suppressed LAP1-induced NE abnormalities. Furthermore, a dominant-negative Torsin induced chromosome segregation defects in a LAP1-dependent manner, suggesting that LAP1 association with chromatin is modulated by Torsin ATPases residing in the perinuclear space and ER lumen. Collectively, our results underscore the importance of dissolving INM protein-chromatin interactions for mitotic fidelity and suggest that LAP1 may not merely be important for Torsin ATPase activation but itself subject to Torsin regulation across the INM.

## Results

### Increased levels of LAP1 impair the dissociation of LAP1–chromatin contacts during mitosis

Although it is generally assumed that changes in posttranslational modifications of INM proteins and chromatin-associated factors, especially protein phosphorylation, induce membrane dissociation from chromatin during mitotic entry, direct evidence in support of this hypothesis in living cells is lacking and our molecular understanding of the process is limited. Here, we set out to gain new insights into the mechanisms required for dissociation of INM proteins from chromatin. In a first step, we tested whether it is possible to impair the release of the NE membranes from mitotic chromatin if any cellular factor involved in dissolving INM protein-chromatin contacts would become limiting by a disturbed ratio between INM proteins and the potential release factor(s). Thus, we overexpressed several abundant INM proteins, including emerin, SUN1, SUN2, LAP2β, LEM2, and LAP1 in HeLa cells. Remarkably, overexpression of LAP1B, the longest human isoform of LAP1, led to severe NE aberrations, while overexpression of other INM proteins did not cause similar defects (Figure 1A). Changes in NE morphology upon LAP1B expression were also evident in other cell types such as HCT116 or HepG2 cells (Figure 1, Supporting Figure 1A). The LAP1B-induced NE aberrations were reminiscent of those that we had previously observed using a synthetic membrane–chromatin tethering system that prevents the release of the NE/ER network from chromatin during mitosis (Champion et al., 2019).

**Figure 1:**
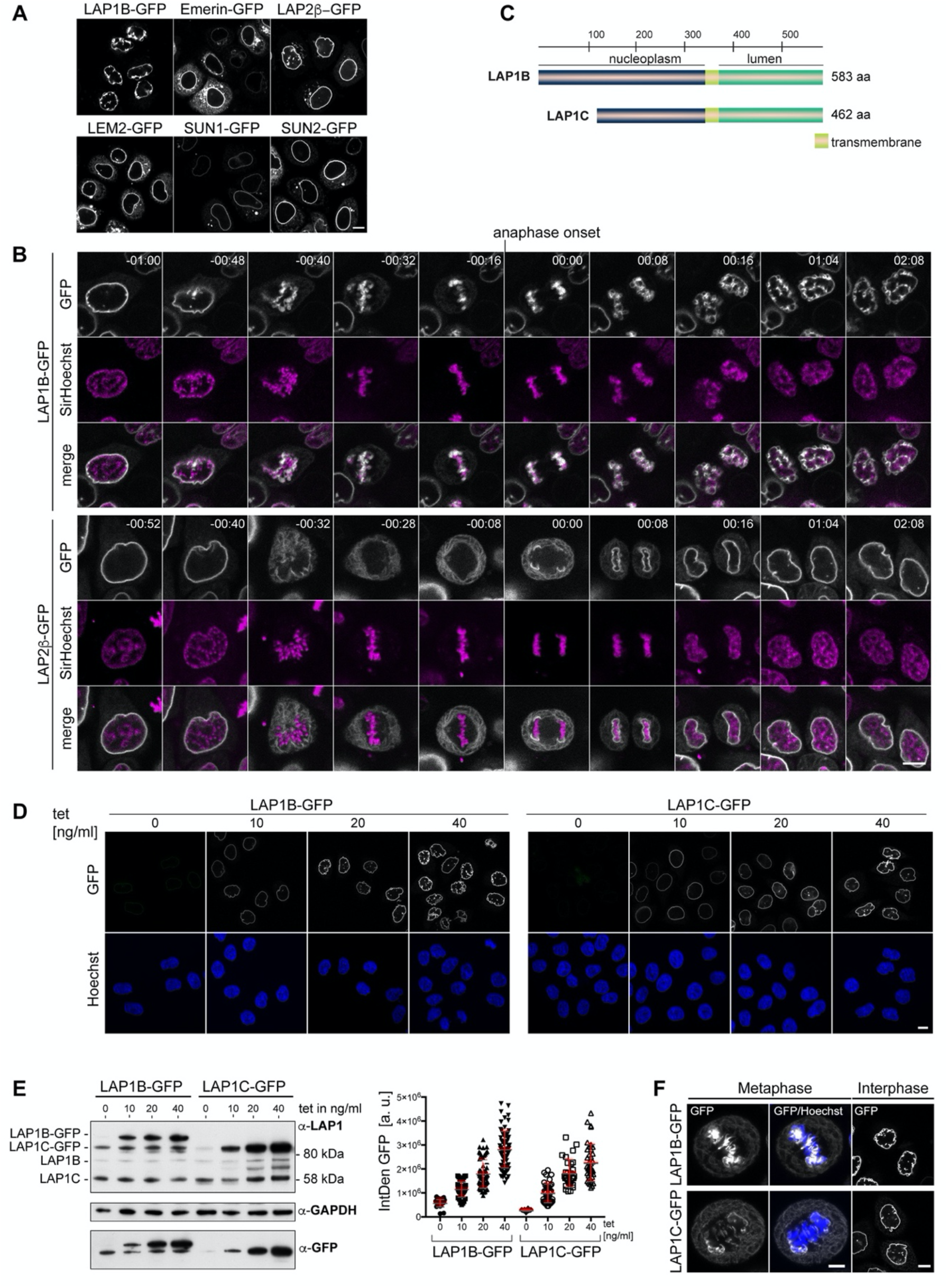
Overexpression of LAP1B and LAP1C causes post-mitotic NE aberrations. (A) Vectors encoding the indicated INM proteins were transfected into HeLa cells. Cells were fixed after 48 h and analyzed by confocal microscopy. Scale bar, 10 μm. (B) Time-lapse images of LAP1B-GFP or LAP2β-GFP expressing HeLa cells progressing through mitosis. DNA was visualized by SirHoechst and used to define anaphase onset (t = 0 min). Scale bar, 5 μm. (C) Scheme depicting the two human LAP1 isoforms, LAP1B and LAP1C. (D) Representative images of stable HeLa cells lines expressing LAP1B-GFP and LAP1C-GFP after induction with different tetracycline (tet) concentrations for 48 h. Scale bar, 10 μm. (E) LAP1 levels were analyzed by immunoblotting of cell lysates (left), or quantification of the integrated GFP density per cell (right). Note that the LAP1B-GFP cell line expressed LAP1C-GFP independent of tetracycline addition, due to an alternative transcriptional start site used for the production of the shorter LAP1 isoform (Santos et al., 2014). (F) Left: Maximum intensity z-projections (5 × 0.63 μm) of confocal images from fixed metaphase HeLa cells expressing LAP1B-GFP or LAP1C-GFP after 48 h of induction. Scale bar, 5 μm. Right: Confocal images of fixed HeLa cells in interphase. Scale bar, 10 μm.

To analyze whether these NE aberrations indeed originated from failed release of LAP1B and thereby membranes from chromatin during mitosis, we performed time-lapse imaging of cells expressing similar levels of either LAP1B-GFP or LAP2β-GFP (Figure 1B, Supporting Figure 1B). Before mitotic entry, nuclei of LAP1B-GFP expressing cells exhibited a normal round shape. Strikingly, throughout mitosis, LAP1B-GFP remained associated with chromatin, and after NE reassembly, both daughter cells exhibited severe nuclear deformations. In contrast, LAP2β-GFP was faithfully released into the mitotic ER, and post-mitotic nuclei of LAP2β-GFP-expressing cells possessed a round unperturbed morphology.

To address how the observed phenotype relates to increasing expression levels of LAP1, we used tetracycline-inducible HeLa cells to tune the amount of LAP1B-GFP. In these experiments, we also included LAP1C, the second isoform of human LAP1 (Figure 1C). LAP1C is produced from a shorter transcript and starts at an alternative translational start site (Met 122), leading to an isoform that is N-terminally truncated by 121 amino residues (Figure 1C) (Santos et al., 2014). Cells were analyzed by confocal microscopy 48 h after tetracycline induction (Figure 1D), and expression levels were examined by quantification of GFP intensity and immunoblotting (Figure 1E). The severity of the NE aberrations indeed increased with rising expression levels for both LAP1B-GFP and LAP1C-GFP (Figure 1D, E), however, they were more pronounced for LAP1B when comparing cells with a similar expression level of both isoforms. We also inspected fixed cells for an association of membranes with chromatin during metaphase (Figure 1F). Both GFP-tagged LAP1 isoforms were enriched at mitotic chromatin, however, the enrichment of LAP1B-GFP was more prominent, indicating a stronger retention of the longer isoform on mitotic chromatin.

To exclude that persistent LAP1–chromatin contacts affect the mitotic localization of endogenous NE proteins, we fixed uninduced and induced LAP1B-GFP cells, and immunostained for nuclear lamins and multiple INM proteins (Figure 1, Supporting Figure 1C). Analysis of metaphase cells revealed that of the tested INM proteins only LAP1B-GFP was strongly enriched at mitotic chromatin. Taken together, overexpression of both human LAP1 isoforms results in their failed removal from mitotic chromatin and cells display an aberrant post-mitotic nuclear morphology, whereas overexpression of other INM proteins at comparable expression levels does not cause similar phenotypes.

### The molecular architecture of the nucleoplasmic domain of LAP1

LAP1 has originally been identified as an INM protein that interacts with nuclear lamins (Foisner & Gerace, 1993, Senior & Gerace, 1988), yet the molecular architecture of its nucleoplasmic domain remains poorly characterized. Fluorescence recovery after photobleaching (FRAP) experiments previously revealed an extremely low diffusional mobility of LAP1B at the INM (Zuleger et al., 2011), but how it is so strongly immobilized at the INM remained to be determined. As overexpressed LAP1B-GFP was associated with chromatin during mitosis, we reasoned that interaction with both chromatin and nuclear lamins may anchor LAP1B at the INM. Our FRAP experiments confirmed that LAP1B-GFP is a highly immobile INM protein (Figure 1, Supporting Figure 2A). In contrast, LAP1C-GFP was more mobile, showing a recovery curve similar to LAP2β-GFP.

**Figure 2.**
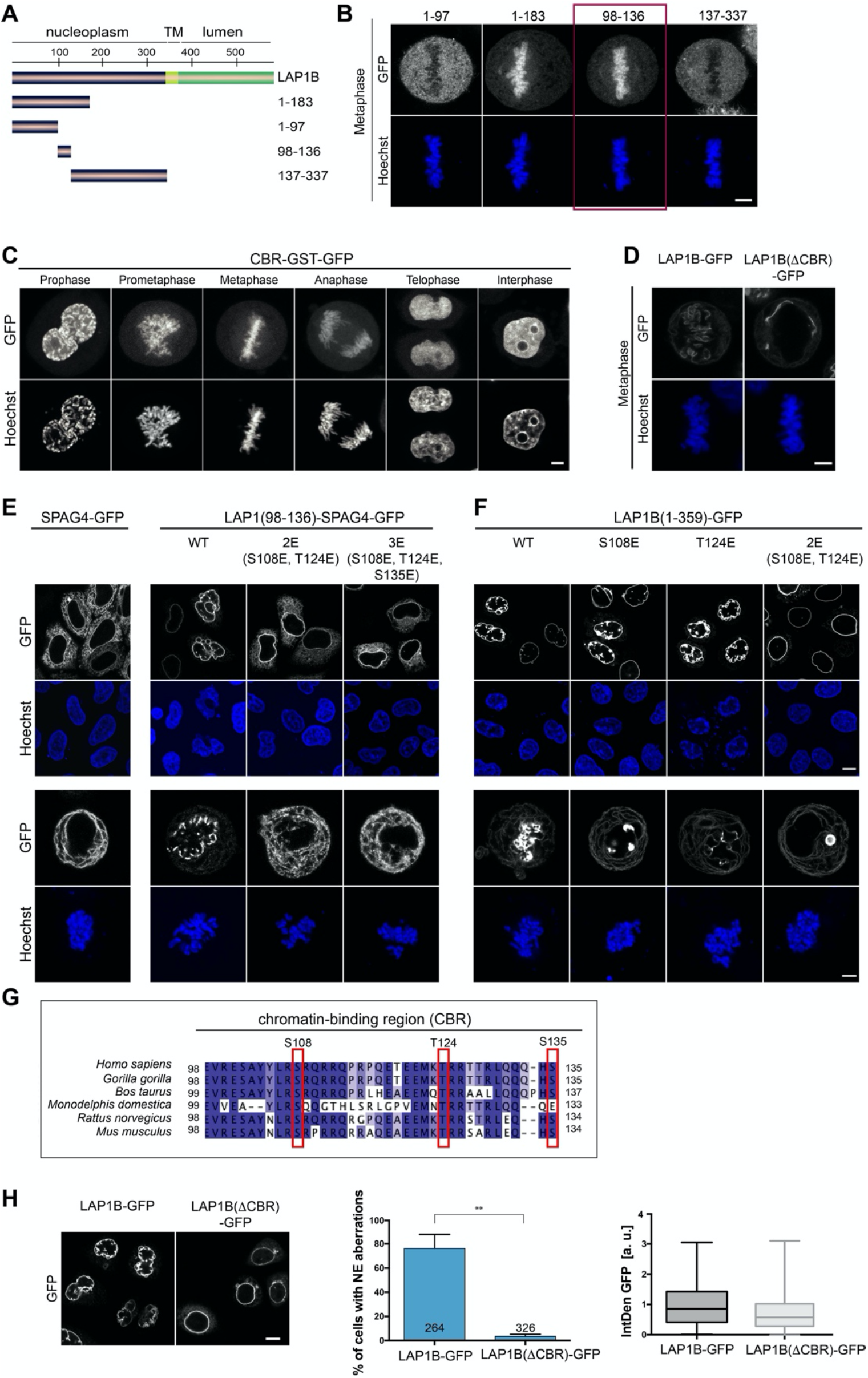
The nucleoplasmic domain of LAP1 contains a central chromatin-binding region that can confer chromatin association during mitosis. (A) Scheme depicting the generated fragments of the nucleoplasmic domain of LAP1B. (B) Metaphase localization of the depicted LAP1 fragments, transiently expressed in synchronized HeLa cells. Maximum intensity z-projections (3 × 1 μm). Scale bar, 5 μm. (C) Localization of LAP1(CBR)-GST-GFP during mitosis in HeLa cells. Scale bar, 5 μm. (D) LAP1B-GFP and LAP1B(ΔCBR)-GFP localization during metaphase. Scale bar, 5 μm. (E) Localization of wild-type SPAG4-GFP and (F) LAP1B(98-136)-SPAG4-GFP derivatives in interphase and prometaphase cells (arrested with nocodazole for 3 h) 48 h after transfection. Scale bars, 5 μm. (F) Alignment of sequences of the chromatin-binding region of LAP1B from the indicated mammalian species. The three mutated residues are boxed in red. (H) Left: Interphase HeLa cells expressing LAP1B-GFP or LAP1B(ΔCBR)-GFP for 48 h. Scale bar, 10 μm. Middle and right: Percentage of nuclei with NE aberrations, quantified in cells expressing LAP1B-GFP or LAP1B(ΔCBR)- GFP at similar levels, based on the integrated GFP density (IntDen) (N=3, n > 264, mean +/- SEM).

To elucidate which lamins contribute to the retention of LAP1 at the NE, we examined the diffusional mobility of LAP1B-GFP and LAP1C-GFP after RNAi-mediated depletion of lamin A/C, lamin B1 or lamin B2 by FRAP (Figure 1, Supporting Figure 2B, C). Downregulation of lamin A/C and lamin B1, but not of lamin B2, led to an increase in the diffusional mobility of LAP1B at the NE. The combined depletion of lamin A/C and lamin B1 had an even stronger effect, indicating that both A- and B-type lamins contribute to retention of LAP1B at the INM. For LAP1C, only the loss of lamin A/C had a prominent effect on its mobility, suggesting that A-type lamins play a major role in retention of the shorter isoform at the INM.

The extraluminal domain of LAP1B is predicted to be largely intrinsically disordered (Figure 2, Supporting Figure 1A), and we set out to delineate its chromatin and lamina-binding regions. By truncation analysis, we loosely delineated two regions in LAP1B that are sufficient for lamina association; an N-terminal fragment of 72 amino acids and a second region comprising residues 184-338 (Figure 2, Supporting Figure 1B-F). To define which part of LAP1 interacts with chromatin, we analyzed the mitotic localization of soluble LAP1B fragments fused to GST-GFP (Figure 2A, B, and Supporting Figure 2A, B). Whereas the N-terminal part (aa 1-183) of LAP1B localized to the metaphase plate, LAP1B(184-337)-GST-GFP was distributed throughout the mitotic cytoplasm. By testing further truncations, we mapped the chromatin-binding region (CBR) of LAP1B to residues 98-136 (Figure 2B). This protein fragment was very efficiently enriched on chromatin throughout mitosis (Figure 2C), and is sufficient for chromatin interaction. Importantly, deletion of the CBR from full-length, membrane-bound LAP1B-GFP abolished its association with chromatin during mitosis (Figure 2D).

Next, we examined whether the CBR contributes to NE localization of LAP1B during interphase. To do so, we chose a domain transfer approach, fusing the CBR of LAP1B to the ER-localized protein SPAG4, as we did before when delineating NE targeting motifs of other INM proteins (Turgay et al., 2010). Consistent with our assumption that chromatin-binding contributes to the retention of LAP1B at the INM, the CBR was sufficient to target the CBR-SPAG4-GFP fusion protein to the NE (Figure 2E). Finally, we tested whether the CBR can interact with chromatin directly by performing electrophoretic mobility shifts assays (EMSAs), revealing that the purified recombinant CBR can interact with reconstituted nucleosomes and even naked DNA (Figure 2, Supporting Figure 2C, D).

Collectively, our mapping experiments delineated three regions within the disordered nucleoplasmic domain of LAP1B that confer interaction with nuclear components: two regions that confer association with the nuclear lamina (aa 1-72 and 184-337) and a central region that is sufficient for chromatin interaction (CBR, aa 98-138; Figure 2G). The multiple regions of LAP1 used for interaction with nuclear partners can very well explain the strong retention of LAP1B at the INM by binding avidity.

### Chromatin-binding region is required for the induction of LAP1-induced NE aberrations

To test whether chromatin interaction of the CBR is required for the post-mitotic NE aberrations induced by LAP1B expression, we analyzed the nuclear shape of cells upon expression of either LAP1B-GFP or LAP1B(ΔCBR)-GFP 48 hours after transfection of the respective constructs. Strikingly, deletion of the CBR abolished the occurrence of NE aberrations (Figure 2H).

To pinpoint residues within the CBR that contribute to mitotic chromatin association, we mutated conserved Ser and Thr residues within the CBR (Figure 2G). With this approach we identified two residues (S108 and T124) that if mutated to glutamate affected NE localization of the CBR-SPAG4-GFP fusion protein during interphase (Figure 2E), abolished its binding to chromatin in metaphase (Figure 2E) and reduced its affinity for DNA and mononucleosomes in vitro (Figure 2, Supporting Figure 2C, D). Although these Ser and Thr residues could in principle be subject to mitotic phosphorylation, we have so far been unable to confirm the potential mitotic phosphorylation of these sites. Comparison of the diffusional mobility of full-length LAP1B-GFP with LAP1B(ΔCBR)-GFP and LAP1B-2E-GFP demonstrated that both mutant constructs exhibited a higher mobility at the NE, highlighting the contribution of the CBR and the identified residues to the retention of LAP1B at the INM (Figure 2, Supporting Figure 3A, B). Importantly, when we introduced both mutations into the membrane-bound, nucleoplasmic domain of LAP1B in the construct LAP1(1-359-2E)- GFP, binding to mitotic chromatin and post-mitotic NE aberrations were fully abolished (Figure 2F). Taken together, this data shows that chromatin-binding contributes to the retention of LAP1 at the NE and identifies two residues (S108 and T124) that are essential for chromatin binding of LAP1B during mitosis. Importantly, this establishes a direct link between the CBR and post-mitotic NE aberrations.

**Figure 3.**
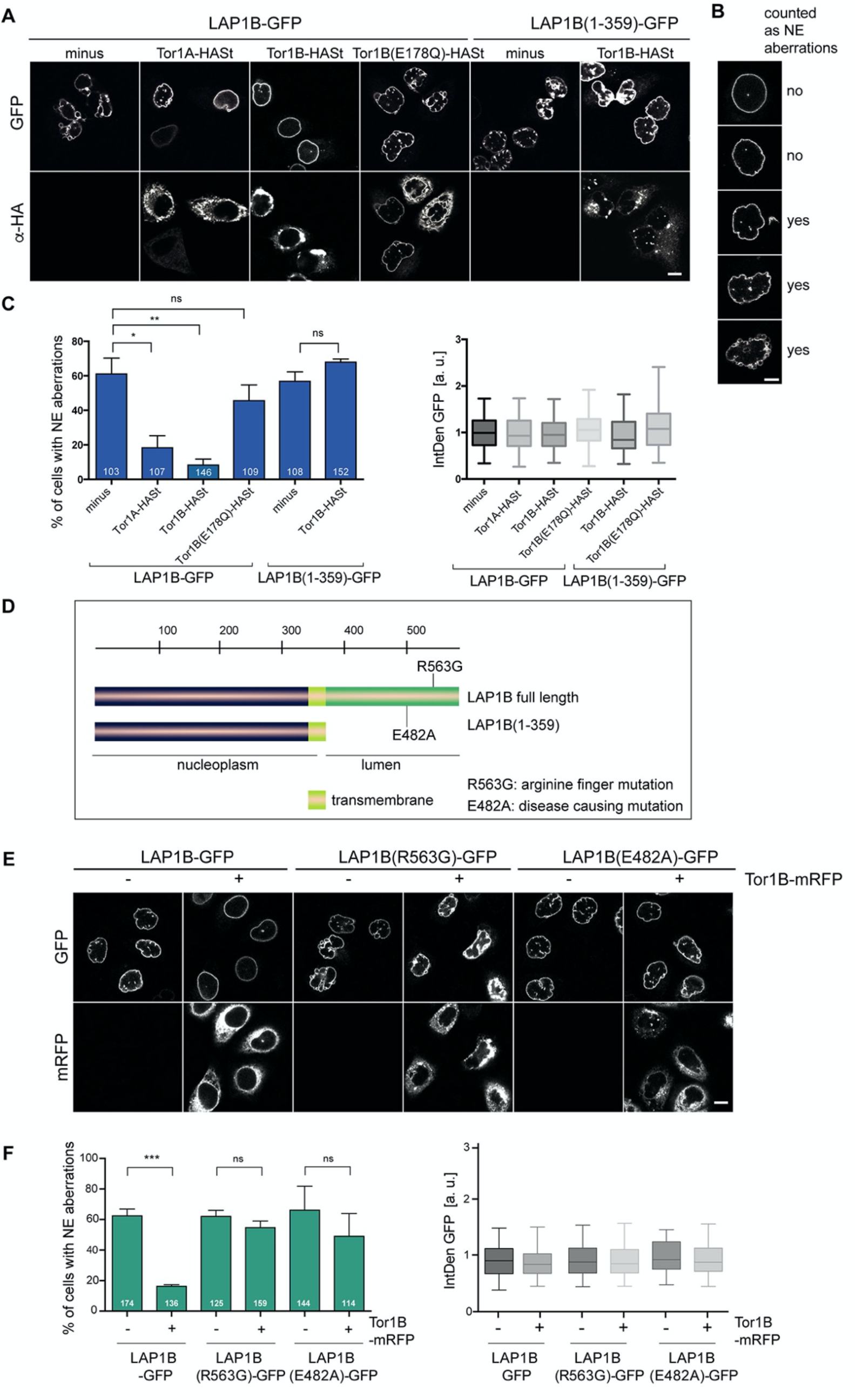
LAP1B-induced NE aberrations can be rescued by co-expression of Torsins. (A) HeLa cells were transiently transfected with constructs encoding for LAP1B-GFP or LAP1B(1-359), either alone or together with the depicted Torsin constructs. After 48 h, cells were fixed, subjected to immunostaining of Torsins using an anti-HA antibody and analyzed by confocal microscopy. Scale bar, 10 μm. (B) Representative images classifying the NE aberration phenotype. Scale bar, 5 μm. (C) The percentage of cells with NE aberrations was quantified (left; N=3, n > 103, mean +/- SEM). Only cells with similar LAP1B-GFP expression levels, based on quantification of the integrated GFP density per cell (IntDen GFP, normalized to the LAP1B minus Torsin control; right) were considered. (D) Depiction of the mutant LAP1B constructs; R563G: Torsin activation deficient-mutant; E482A, disease-causing mutant. (E) HeLa cells were transiently transfected with constructs encoding for the indicated LAP1B-GFP derivatives, either alone or together with Tor1B-mRFP. After 48 h, cells were fixed and analyzed by confocal microscopy. Scale bar, 10 μm. (F) The fraction of cells with NE aberrations was quantified as in D (N=3, n > 114, mean +/- SEM).

### LAP1B-induced nuclear envelope aberrations can be rescued by Torsin co-expression

LAP1 is an established activator of Torsin AAA+ ATPase family members, i.e. of Torsin1A (Tor1A) and Torsin1B (Tor1B) (Brown et al., 2014, Sosa et al., 2014). The persistent chromatin association of overexpressed LAP1B during mitosis prompted us to examine whether restoring a normal LAP1B to Torsin ratio by co-overexpression of these proteins would rescue NE shape defects. We transiently expressed LAP1B-GFP alone or together with either Tor1A or Tor1B, both C-terminally tagged with an HA-Strep tag, and analyzed the percentage of cells containing NE aberrations 2 days after transfection. For quantification, we binned cells into a window of comparable fluorescence intensities to exclude an influence of LAP1B expression levels. Indeed, co-expression of Torsins led to a striking decrease in NE aberrations (Figure 3A, B, C).

To test whether this rescue depends on the ATPase activity of Torsins, we utilized a well-characterized Glu to Gln mutation in the Walker B motif that prevents ATP hydrolysis but not ATP binding, and can be used to lock AAA+ ATPases on their substrates (Snider et al., 2008, Weibezahn et al., 2003). In the case of Torsins, the ATPase deficient E to Q mutation traps Torsins on its activators LAP1 and LULL1 (an ER-resident LAP1 paralog), while substrates of Torsins have not been identified (Jungwirth et al., 2010, Rose et al., 2014, Zhao et al., 2013). In contrast to wild-type Torsins, Tor1B(E178Q)-HASt overexpression failed to rescue the NE aberrations caused by LAP1B-GFP, indicating that the enzymatic activity of Torsins is needed to prevent the LAP1B-induced phenotypes and not the LAP1B to Torsin ratio alone (Figure 3A, B, C). As expected, NE aberrations triggered by expression of LAP1B(1-359)-GFP, which lacks the luminal domain required for Torsin activation, were not rescued by Tor1B-HASt overexpression (Figure 3A, B, C). Similarly, Tor1B co-expression did not ameliorate NE aberrations caused by expression of LAP1B(R563G) (Figure 3E, F), a mutant inactivating the Arg-finger needed for stimulation of Torsin ATPase activity (Brown et al., 2014, Sosa et al., 2014). Another non-functional LAP1B mutant, E482A, known to cause severe dystonia, cerebellar atrophy and cardiomyopathy (Dorboz et al., 2014), also induced NE aberrations, which were not prevented by Torsin co-expression. Thus, Torsins can only act on LAP1 derivatives that are competent for the activation of its ATPase activity.

We also examined the influence of the ER-localized Torsin activator LULL1 as it had been suggested to regulate access of Torsins to the INM (Goodchild et al., 2015, Vander Heyden et al., 2009). However, expression of LULL1 did not improve the NE shape of LAP1B-GFP expressing cells (Figure 3, Supporting Figure 1). Taken together, the correct LAP1B to Torsin ratio and the functional interplay between LAP1B and Torsins ensure an unperturbed post-mitotic nuclear morphology.

### Torsins influence the diffusional mobility of LAP1 at the NE

Given that Torsins are expressed in postmitotic, differentiated cells, we next set out to examine whether Torsins influence LAP1 dynamics at the NE during interphase. Firstly, we compared the mobility of the Torsin activation-deficient mutant R563G and the disease-causing mutant E482A to wild-type LAP1B by FRAP. The E482A mutation has been suggested to impair folding of the luminal domain (Demircioglu et al., 2016). Since LAP1B is so strongly immobilized at the INM by binding to nuclear lamins and to chromatin, it was not surprising that no mobility changes were detected (Figure 4A). However, upon depletion of laminA/C and laminB1, it became apparent that both mutants were less mobile compared to the wild-type protein (Figure 4B). Secondly, to more directly test whether Torsins might influence the retention of LAP1 at the INM, we measured the mobility of LAP1B in Tor1B wild-type and Tor1B(E178Q)-expressing cells in the presence and absence of nuclear lamins (Figure 4C, D). The presence of the dominant-negative Torsin indeed significantly reduced the mobility of LAP1B-GFP in laminA/C and B1-depleted cells. Thirdly, to address how chromatin interaction by the CBR of LAP1B influences this behavior, we exploited the identified 2E mutant that reduces the affinity of the CBR for chromatin (Figure 2, Supporting Figure 2C, D). As expected, the LAP1-2E mutant was more mobile, and its mobility seemed less reduced by Tor1B(E178Q) expression.

**Figure 4:**
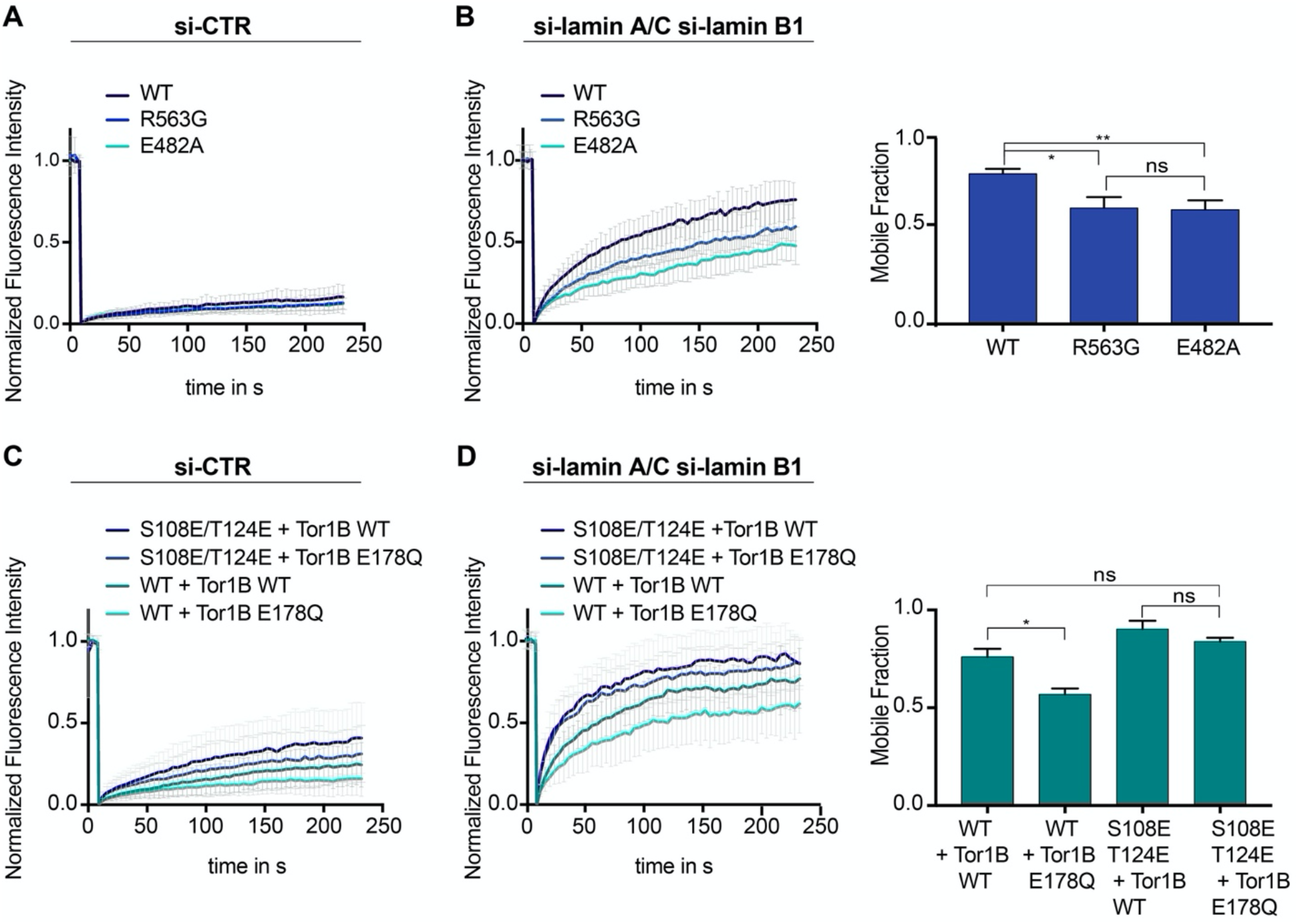
Mobility of LAP1B at the NE of interphase cells is influenced by Torsins. (A) FRAP analysis of HeLa cells expressing either wildtype LAP1B-GFP or the indicated mutant variants and treated with a control siRNA (N=5, n >18, mean +/- SD). (B) FRAP analysis as in B after RNAi-mediated co-depletion lamin A/C and lamin B1 (N = 5, n > 21, mean +/- SD). Corresponding mobile fractions (mean +/- SEM, *p<0.05). (C) FRAP analysis of LAP1B-GFP and the chromatin-binding deficient variants in cells expressing either Tor1B wild-type or the dominant-negative Tor1B(E178Q) mutant both tagged with mRFP, and treated with a control siRNA (N = 4, n >18, mean +/- SD). (D) FRAP analysis in lamin-depleted cells as in C (N = 4, n > 25, mean +/- SD). Corresponding mobile fractions (mean +/-, *p<0.05).

### LAP1 mutants unable to activate Torsins cause binucleation that depends on LAP1–chromatin interaction

We have previously shown that persistent membrane–chromatin interactions during mitosis leads to severe post-mitotic NE morphology defects, and, in addition, to a significant increase in chromosome segregation defects, eventually resulting in cell division failure and an increase in the number of binucleated cells (Champion et al., 2019). To elucidate whether persistent LAP1–chromatin interactions would mimic these phenotypes, we examined binucleation upon expression of LAP1B-GFP and its derivatives deficient in Torsin activation. To quantify binucleated cells, we visualized the cytoplasmic contours by tubulin immunofluorescence and counterstained nuclei with Hoechst (Figure 5A, B, C). Compared to the parental HeLa cell line, expression of LAP1B-GFP induced a two-fold increase in binucleated cells. At a similar level, both LAP1B(1-359)-GFP and LAP1B(R563G)-GFP led to a twofold further rise in the number of binucleated cells compared to cells expressing LAP1B wild-type. This number was even higher for the disease-causing mutant LAP1B(E482A)-GFP. Thus, the E482A mutation seems to be a very potent dominant-negative if expressed.

**Figure 5.**
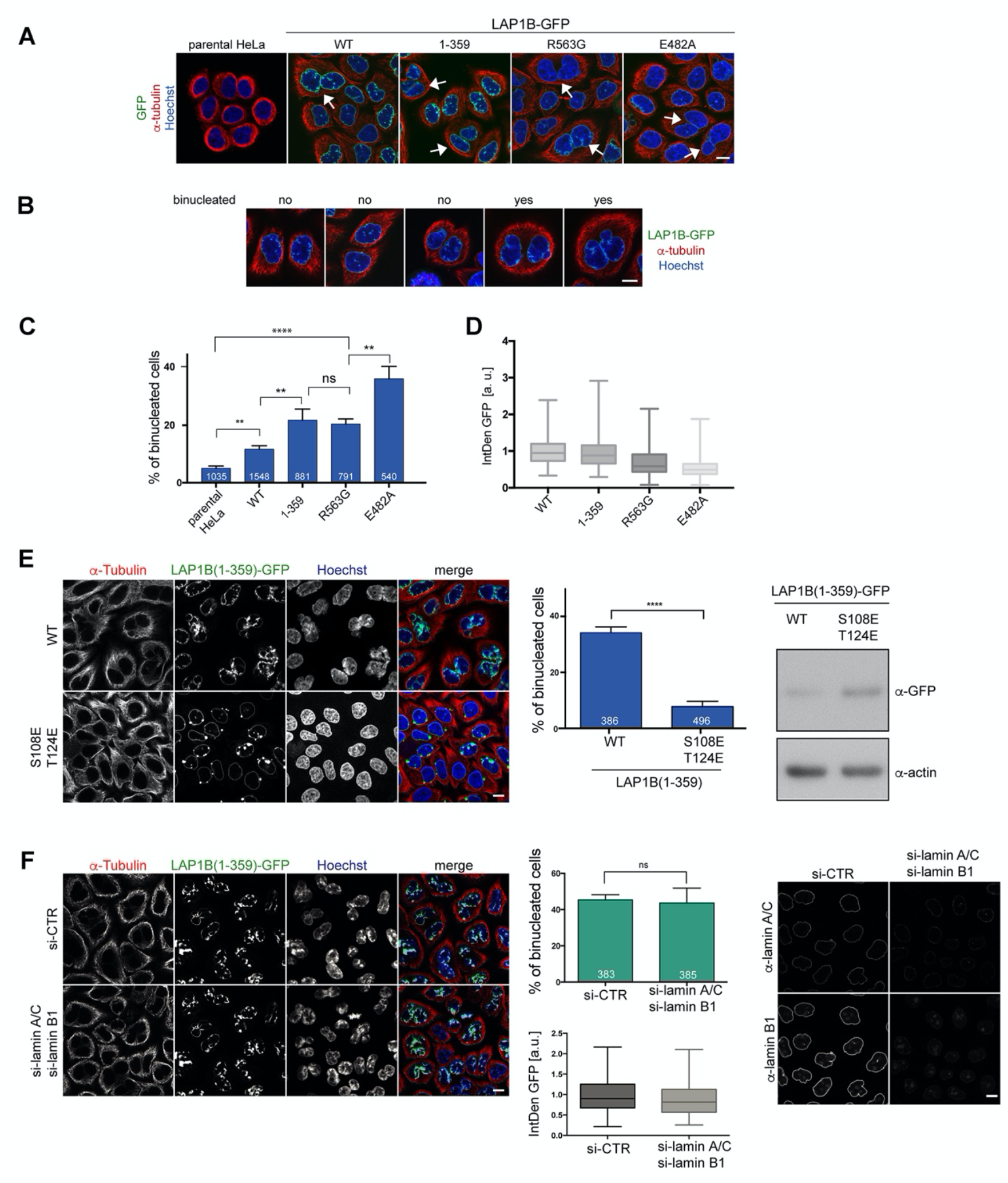
LAP1B mutants deficient in Torsin activation lead to increased binucleation dependent on chromatin interaction of LAP1. (A) Expression of LAP1B(1-359)-GFP was induced with 0.01 μg/ml tetracycline and expression of LAP1B-GFP wild-type, LAP1(R563G)-GFP and LAP1(E482A)-GFP with 1 μg/ml tetracycline for 48 h. The parental HeLa cell line was used as a negative control. Then, cells were fixed, immunostained for tubulin and analyzed by confocal microscopy. Scale bar, 10 μm. White arrows denote binucleated cells. (B) Images representing the categories used for classification of binucleation. (C) Quantification of the fraction of binucleated cells (N = 3, n > 540, mean +/-SEM). (D) The integrated density (IntDen) of the GFP signal was measured and normalized to LAP1B-GFP. (E) Binucleation caused by LAP1B is prevented by point mutations that abolish chromatin-binding. Expression of LAP1B(1-359)-GFP or LAP1B(1-359-2E)-GFP was induced with tetracycline (0.01 μg/ml and 1 μg/ml, respectively) in stable HeLa cell lines for 48 h. Cells were immunostained for tubulin and counterstained with Hoechst. Binucleated cells were quantified (N = 3, n > 300, mean +/- SEM; ****p<0.0001). Expression levels of LAP1B(1-359)-GFP and LAP1B(1-359-2E)-GFP were compared by immunoblotting. (F) LAP1B(1-359)-GFP expression was induced for 48 h as in E in either mock-treated or in lamin A/C and lamin B1-co-depleted cells (RNAi for 72 h, induction of LAP1B constructs for the last 48 h). Cells fixed and analyzed as in E. The number of binucleated cells was quantified (N = 3, n > 383, mean +/- SEM). LAP1B(1-359)-GFP expression levels were determined based on the integrated GFP intensity (IntDen). Lamin RNAi was controlled by immunofluorescence. Scale bars, 10 μm.

Next, we addressed the contribution of chromatin interaction of LAP1 to the formation of binucleated cells by comparing HeLa cell lines expressing either LAP1B(1-359)- GFP or the chromatin-binding deficient 2E (S108E, T124E) mutant, comparing cells at a high expression level by using an elevated tetracycline concentration for induction of the constructs. Whereas LAP1B(1-359)-GFP caused binucleation in nearly 40% of cells under these conditions, LAP1B(1-359-2E)-GFP did not (Figure 5E, F), confirming that binucleation is caused by the LAP1B-chromatin interaction. As LAP1B also interacts with lamin A/C and lamin B1, we further tested whether LAP1–lamina interactions contribute to binucleation. As RNAi-mediated depletion of lamin A/C and laminB1 in cells expressing LAP1B(1-359)-GFP did not reduce the number of binucleated cells (Figure 5G, H), we conclude that the cell division defects are independent of nuclear lamins, and thus solely related to the aberrant binding of LAP1 to chromatin during mitosis.

### An ATPase-deficient Torsin impairs the removal of endogenous LAP1 from mitotic chromatin and increases binucleation

To date, five human Torsins have been identified. Because these five members of the Torsin family likely function, at least in part, in a redundant manner, it is challenging to reach a sufficient downregulation of these enzymes by conventional protein depletion experiments. As dominant-negative approaches have been decisive in elucidating the cellular function of ATPases, we therefore decided to investigate the consequences of expression of the dominant-negative Tor1B(E178Q) mutant. Strikingly, when we visualized endogenous LAP1 by immunofluorescence in cells expressing Tor1B(E178Q)-GFP, we observed LAP1-positive patches localizing to (pro)- metaphase chromatin, indicating an impaired release of LAP1 from chromatin during mitotic entry (Figure 6A). In comparison, cells expressing wild-type Tor1B-GFP showed GFP-positive membrane patches in the peripheral mitotic ER. These patches are reminiscent of previously described ER membrane foci in TorsinB-expressing interphase cells (Rose et al., 2014).

**Figure 6.**
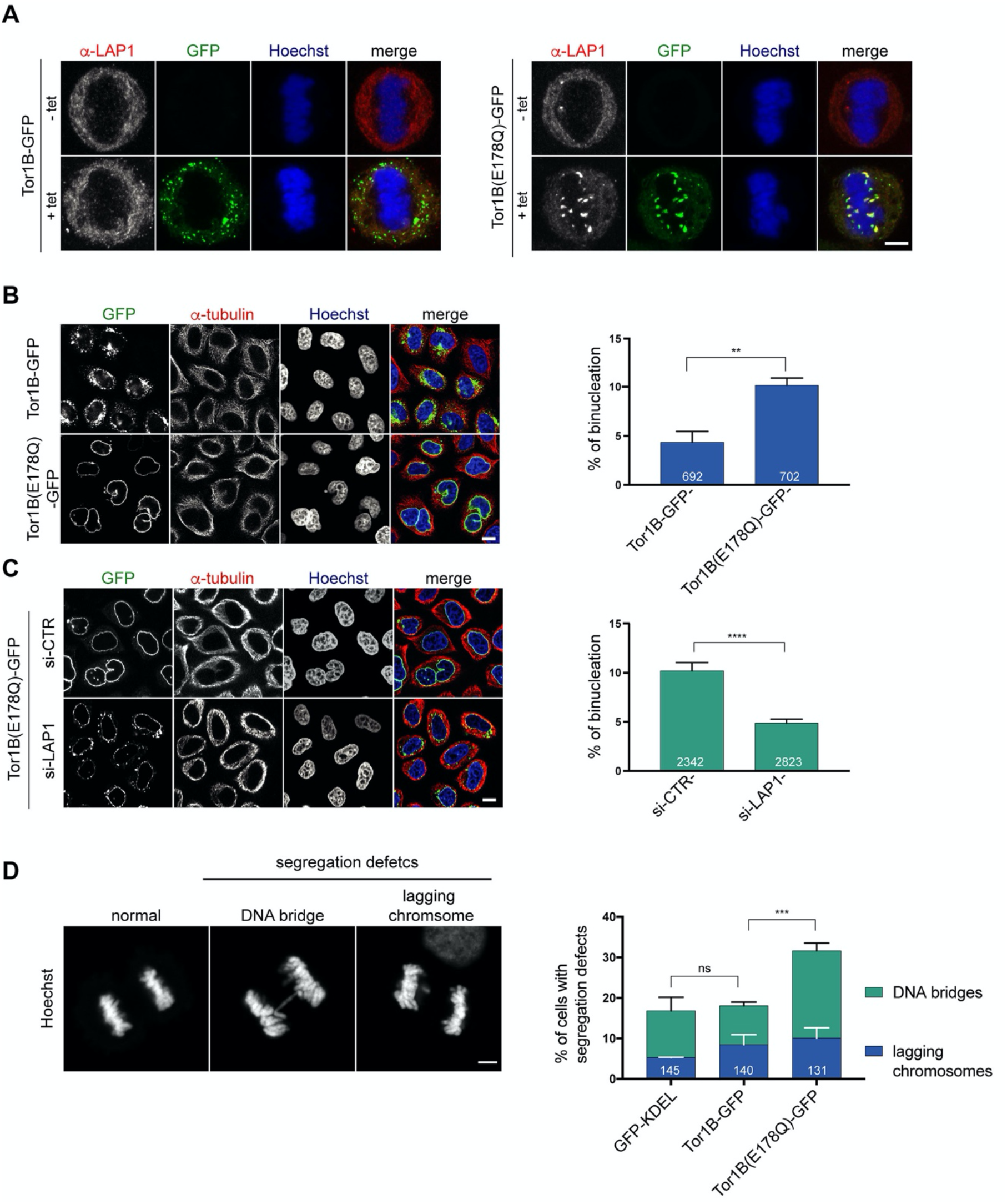
A dominant-negative, ATPase-deficient Torsin1B mutant increases binucleation in a LAP1-dependent manner. (A) Endogenous LAP1 remains associated with chromatin in metaphase upon overexpression of Tor1B(E178Q)- GFP. Representative maximum intensity projections (5 x 0.63 μm) of confocal images of metaphase HeLa cells upon induction of either Tor1B-GFP and Tor1B(E178Q)-GFP expression with 1 μg/ μl tetracycline for 48 h. Localization of endogenous LAP1 was analyzed by immunofluorescence staining. Scale bar, 5 μm. (B) Confocal images of Tor1B-GFP and Tor1B(E178Q)-GFP-expressing cells stained with Hoechst and immunostained for tubulin. Scale bar, 10 μm. Quantification of binucleated cells (N =3, n > 835, mean +/- SD). (C) Expression of Tor1B(E178Q)-GFP was induced for 48 h in LAP1-depleted or mock-treated cells. Binucleation was analyzed by confocal microscopy and quantified (N= 3, n > 1915, mean+/- SEM). Scale bar, 10 μm. (D) Quantification of chromosome segregation defects (DNA bridges and lagging chromosomes) in fixed GFP-KDEL, Tor1B-GFP and Tor1B(E178Q)-GFP expressing anaphase cells after 24 h of induction with tetracycline. Segregation errors were manually quantified (N = 3, n > 130, mean+/- SEM).

Next, we compared the formation of binucleated cells upon induction of Tor1B(E178Q)- GFP or wild-type Tor1B-GFP. Remarkably, Tor1B(E178Q)-GFP expression led to a twofold higher number of binucleated cells as compared to cells expressing Tor1B-GFP wild-type (Figure 6B). Notably, no post-mitotic NE aberrations were observed in Tor1B(E178Q) expressing cells, which can be explained by the lack of persistent LAP1/Tor1B-E178Q-GFP chromatin interactions in anaphase/telophase.

As binucleation is indicative for a failed cytokinesis and commonly caused by chromosome segregation defects, we quantified DNA bridges and lagging chromosomes. This analysis revealed a significant increase in chromosome segregation defects (Figure 6D). To elucidate whether the observed increase in binucleated cells can be attributed to the failed removal of LAP1 from mitotic chromatin, we performed RNAi-mediated depletion of LAP1 in Tor1B(E178Q)-GFP expressing cells. Importantly, the number of binucleated cells was significantly reduced in LAP1-depleted cells (Figure 6C). As judged by the GFP signal, the Tor1B(E178Q)-GFP expression seemed slightly reduced in LAP1-depleted cells as compared to cells treated with control siRNA (Figure 6C; Figure 6, Supporting Figure 2), and we wished to exclude that unequal Tor1B(E178Q) expression levels led to the reduction of binucleation in LAP1-depleted cells. Therefore, we induced Tor1B(E178Q) with a lower tetracycline concentration in control cells. Even though these cells now expressed two-fold less Tor1B-E178Q compared to LAP1-depleted cells, binucleation was still significantly reduced upon downregulation of LAP1 (Figure 6, Supporting Figure 2A, B, C). For comparison, we also depleted the ER-resident Torsin-activator LULL1 in Tor1B(E178Q)-GFP expressing cells, but could not observe a reduction in binucleated cells (Figure 6, Supporting Figure 1C, D). LULL1 depletion also did also not prevent the localization of LAP1 to mitotic chromatin upon Tor1B(E178Q) expression (Figure 6, Supporting Figure 1E). Taken together, the dominant-negative Tor1B variant induces the persistence of LAP1-positive membrane patches on (pro)-metaphase chromatin, chromosome segregation defects and binucleation in a LAP1-dependent manner. Collectively, our findings indicate that Torsin function is important for mitotic fidelity by assuring normal LAP1 behavior during mitosis.

## Discussion

During NE breakdown, membrane proteins of the INM are released from chromatin and the nuclear lamina to allow for the spatial separation of membranes from chromatin. Our work uncovers that the abundant and ubiquitously expressed INM protein LAP1 is tightly anchored at the INM of interphase cells by association with both chromatin and the nuclear lamina. Unexpectedly, we found that LAP1 remains stuck on chromatin in mitosis if either the stoichiometry of LAP1 to Torsins is unbalanced or Torsin functionality is compromised. These findings are intriguing for two reasons. Firstly, they indicate that Torsins, residing in the perinuclear space, influence LAP1’s ability to bind chromatin at the other side of the membrane. Secondly, reminiscent of some of the phenotypes that we have previously observed using a synthetic membrane-chromatin tether (Champion et al., 2019), the failure in release of LAP1 from mitotic chromatin is associated with chromosome segregation defects, binucleation and postmitotic NE aberrations.

### The Torsin-LAP1 interplay

Based on studies in human cells, mice and Drosophila, diverse roles of Torsins have been suggested, including functions in ER homeostasis, secretion, lipid metabolism, NPC biogenesis, and the nuclear export of large particles like mRNPs or HSV1 capsids by a vesicular transport process across the NE (Chen et al., 2010, Grillet et al., 2016, Jokhi et al., 2013, Maric et al., 2011, Nery et al., 2011, Pappas et al., 2018, Rampello et al., 2019, Shin et al., 2019, Turner et al., 2015). However, while all these suggested functions are supported by the reported cellular phenotypes, biochemical and genetic data, it is currently unclear what are direct or indirect consequences of Torsin function since the molecular mechanism of Torsin action has remained elusive. All attempts to identify bona fide substrates of these unsual, ER-luminal AAA+ ATPases have so far been unsuccessful. It has therefore even been speculated that the established Torsin activators LAP1 and LULL1 could act simultaneously as activators and substrates of these mysterious ATPases (Cascalho et al., 2017, Rose et al., 2015), especially since LAP1 and LULL1 are highly enriched on the ATP-bound form of some Torsins, as expected for substrates of AAA+ ATPases.

So, how do our findings relate to the current understanding of the molecular function of Torsins? Our experiments suggest that the interaction of Torsins with the C-terminal, luminal domain of LAP1 in the ER lumen influences the chromatin-binding activity of LAP1’s nucleoplasmic domain on the other side of the membrane. This implies that LAP1 might not merely behave as an activator of Torsins, but could in turn itself be a downstream effector or ‘substrate’ influenced by the activity of the ATPases. Two observations indicate that this reverse crosstalk from Torsin to LAP1 is indeed linked to the enzymatic activity of the ATPase: Firstly, LAP1B-induced NE aberrations can be rescued by expression of TOR1B wild-type but not by an ATPase inactive mutant. Secondly, LAP1 derivatives that are deficient in Torsin ATPase activation execute a stronger dominant-negative effect on cell division than wild-type LAP1.

Clearly, even if considering LAP1 a ‘substrate’ of Torsins ATPases, their interplay must differ from the ‘classical’ enzyme-substrate mechanism typical of AAA+ ATPases that thread protein substrates through the central pore of a usually hexameric ATPase rings (Hanson & Whiteheart, 2005, Olivares et al., 2016). Although Torsins show sequence similarity to bacterial ClpX ATPases that serve as the paradigm of substrate-threading AAA+ ATPases (Olivares et al., 2016), they seem to lack the characteristic pore loops involved in substrate gripping during the translocation of an unfolded protein chain through the central opening of the ring (Demircioglu et al., 2016). Furthermore, the luminal domains of LAP1 and LULL1 are structural mimics of the ATPase fold. These domains bind laterally to the ATPase domain of Torsins to activate ATP hydrolysis through their arginine finger (Brown et al., 2014, Sosa et al., 2014), inconsistent with LAP1 being a substrate of a threading mechanism. Consistently, Torsins have not been found to unfold the luminal domains of LAP1 and LULL1 in vitro (Zhao et al., 2013). Thus, the interplay of Torsin and LAP1 must work differently.

From a conceptual point of view, we can envisage several scenarios as to how Torsin ATPase activity could influence chromatin interaction of LAP1. One possibility is that activation of Torsins transduces mechanical forces across the membranes, thereby weakening the interaction of LAP1 with chromatin, and thereby facilitating its removal from chromatin. This might entail a conformational rearrangement of the luminal domain of LAP1 relative to the ATPase domain of Torsins induced by ATP hydrolysis, and propagated to the other side of the membrane either in form of a Torsin-induced pulling on LAP1 or a rotational movement of LAP1 along its longitudinal axis in the plane of the membrane. Another plausible scenario is that Torsin ATPase activation leads to the disassembly of LAP1 dimers or higher order oligomers by affecting their lateral association, thereby influencing the avidity of the LAP1–chromatin interaction.

Based on structural studies that revealed an AAA+ fold mimicry of the luminal domains of LAP1 and LULL1, it has originally been assumed that they form mixed heterohexameric rings with Torsins, with support from in vitro assembly experiments (Brown et al., 2014, Sosa et al., 2014). Such a heterohexameric configuration with 3 LAP1 and 3 Torsin molecules per hexamer would readily explain how an ATPase-deficient Torsin mutant such as Tor1B(E178Q) could increase the affinity of LAP1 for chromatin due to avidity effects caused by the entrapment of LAP1 in these ATP-stabilized ring structures.

However, it was subsequently noted that the surfaces of the luminal domains of LAP1 and LULL1 are only well conserved on their catalytic interface with Torsins but not on the potential non-catalytic interface, indicating the lack of evolutionary pressure to maintain this interface and shedding some doubt on the suggested hetero-hexameric organization (Chase et al., 2017, Demircioglu et al., 2016). In contrast, Torsins display surface conservation on both their catalytic and non-catalytic interfaces, consistent with the possibility that Torsins could form homo-oligomers. Indeed, Torsins can assemble into ATP-dependent, homo-oligomeric structures in vitro, adopting the configuration of a spiral or potentially a lock-washer (Chase et al., 2017, Demircioglu et al., 2016). Such configuration would provide only one accessible interface for binding of LAP1 or LULL1 to promote oligomer disassembly upon ATPase activation. It remains to be seen which higher order configuration of Torsins exist in living cells, also in light of the view that LAP1 and LULL1 are overstoichiometric to Torsins (Itzhak et al., 2016). Such knowledge will be key to inform rational models of how Torsins influence chromatin interaction of LAP1.

### LAP1 interaction with chromatin and the nuclear lamina

LAP1B possesses an extremely low diffusional mobility in comparison to other INM proteins (Zuleger et al., 2011). Its intrinsically disordered nucleoplasmic domain engages in interaction with at least three binding partners that contribute to its strong retention at the INM, namely lamin A/C, lamin B1 and chromatin. This raises the question of whether the strong immobilization of LAP1 at the NE might bear an additional function beyond retaining the protein at the INM. Potential roles include the mechanical coupling of the INM to chromatin and the nuclear lamina or a function in chromatin positioning at the nuclear periphery, e.g. in heterochromatin organization. INM proteins together with nuclear lamins are known to serve as heterochromatin tethers that promote the positioning of transcriptionally silenced heterochromatin domains underneath the nuclear envelope of differentiated cells (Guelen et al., 2008, Solovei et al., 2013). Considering the organization of its nucleoplasmic domain, LAP1 seems ideally suited to bridge lamina-chromatin interactions. Notably, LAP1 is developmentally essential as LAP1 knockout mice exhibit perinatal lethality (Kim et al., 2010). Recently, loss of LAP1 expression in human patients caused by a mutation leading to an in-frame termination codon in LAP1 has been associated with an early onset multisystemic nuclear envelopathy manifesting in severe neurological and developmental deficiencies, resulting in early lethality of the patients (Fichtman et al., 2019). These severe defects are most likely caused by a lack of Torsin activation at the NE by lack of LAP1. However, it might also be worth considering whether LAP1 might have a chromatin-associated function that is needed for proper development. Our study primarily focused on the long isoform LAP1B. In comparison, LAP1C is predicted to possess a shortened chromatin-binding region. When overexpressed, LAP1C is also capable of binding to chromatin during mitosis, albeit to a lesser extent than LAP1B. Interestingly, the relative expression of LAP1B and LAP1C differs between tissues and changes during development (Santos et al., 2014, Shin et al., 2014). While LAP1C is the predominant isoform in undifferentiated cells, the relative levels of LAP1B strongly increase during differentiation and, interestingly, upon induced cell cycle exit of cultured somatic cells (Santos et al., 2015). In light of our observation that LAP1B is more prone to stick to mitotic chromatin than LAP1C and induces mitotic errors when present at increased levels in rapidly dividing cells, the physiological upregulation of LAP1B in correlation with cell cycle exit seems ideally suited to prevent potentially fatal mitotic defects caused by LAP1B.

### The gist of the matter – mitosis or interphase?

Although we present evidence that Torsins influence the interaction of LAP1 with chromatin, we don not necessarily assume that Torsins have evolved to liberate LAP1 from chromatin for open mitosis or that they possess a function dedicated to mitosis. Given that primary dystonia caused by mutations of Tor1A or LAP1 seems to arise through a dysfunction of post-mitotic neurons, the function of Torsins is probably essential for the homeostasis of interphase cells. We rather consider it likely that mitosis is perturbed as a consequence of a perturbed LAP1-Torsin interplay during interphase that manifests in the subsequent mitosis.

Both the expression of the Tor1B ATPase-deficient E178Q mutant or excess LAP1 impaired the full removal of LAP1 from chromatin in mitosis, and is associated with chromosome segregation errors, increased binucleation of cells and post-mitotic NE aberrations. When we weakened chromatin association of LAP1B by mutation of two residues in the chromatin-binding region to phospho-mimetic residues, these phenotypes disappeared, suggesting that the prolonged chromatin association of LAP1 during mitosis is the underlying reason. Dissociation of membrane chromatin contacts is currently assumed to be primarily controlled by protein phosphorylation. While our own investigations of mitotic phosphorylation of these sites remained inconclusive, one of these residues, i.e. Ser135, has indeed been detected as mitotically phosphorylated in large scale phosphoproteomics (Dephoure et al., 2008). If chromatin interaction of LAP1 were normally controlled by (mitotic) kinases, then the perturbed Torsin-LAP1 interplay could also affect the access of kinases to the nucleoplasmic domain of LAP1, e.g. by a failure in dissolving interactions of LAP1 with either itself or other partners. Interestingly, the nucleoplasmic domain of LAP1 is known to interact with protein phosphatase 1, the persistent binding of which could counteract the action of (mitotic) kinases (Santos et al., 2013).

Both binucleation and aneuploidy may contribute to or even drive tumorigenesis (Chow et al., 2012, Levine & Holland, 2018, Tanaka et al., 2018). Because Torsins likely function in a redundant manner, it is highly improbable that Torsin function can be completely abolished by mutations in most tissues. For LAP1, in contrast, mutations that specifically impair its Torsin binding or activation capability, present a risk of causing mitotic errors. Remarkably, mutation of LAP1’s arginine finger was identified as a somatic mutation in melanoma (Krauthammer et al., 2012) and colon cancer (Cancer Genome Atlas, 2012). However, whether this mutation genuinely contributes to tumorigenesis remains to be clarified. Based on our findings, one could expect aneuploidy to develop over time if differentiated cells with high LAP1B expression levels were forced to reenter the cell cycle for instance by oncogenic driver mutations.

## Material and Methods

### Molecular cloning

The LAP1B, LULL1, Tor1A and Tor1B coding regions were amplified from HeLa cDNA and cloned into the vector pcDNA5.1-GFP FRT/TO (cytomegalovirus [CMV] promoter; Invitrogen) for generation of stable, tetracycline-inducible cell lines. The respective constructs for LAP1B (isoform 3) and Tor1B pcDNA5.1 FRT/TO served as PCR or mutagenesis templates, yielding the following derivatives: LAP1C, LAP1B(1-359), LAP1B(Δ98-136), LAP1B(73-end), LAP1B(184-end), LAP1B(R563E), LAP1B(E482A), LAP1B(S108E,T124E), LAP1B(1-359)(S108E,T124E), LAP1B(S108E,T124E, R563G), LAP1(98-136), LAP1(98-136)(S108E,T124E) and Tor1B(E178Q). Fragments of the nucleoplasmic domain of LAP1 were subcloned into the mammalian expression vector pK7-GST-GFP (Ungricht et al., 2015) and/or pK7-GST-GFP-NLS(SV40) (Erkmann et al., 2005) or the bacterial expression vector pQE60-zz-6xhis (Kutay et al., 1997), encoding for two z domains derived from the immunoglobulin-binding domain of *Staphylococcus aureus* Protein A. For the domain swap experiment with SPAG4, the chromatin binding region of LAP1 (aa 98-136) and mutants thereof were cloned into pEGFP-N3-SPAG4 (Turgay et al., 2010). The emerin and LEM2 coding regions were amplified from HeLa cells derived cDNA and cloned into pEGFP-N3 (Clontech). pEGFP-N3-derived plasmids encoding for LAP2β (Ungricht et al., 2015), SUN1 and SUN2 (Turgay et al., 2010) have been described before. Point mutations were introduced by using QuikChange site-directed mutagenesis.

### Antibodies

The following commercial antibodies were used in this study: anti-β-actin (mouse; Sigma, A1978; RRID: AB_476692), anti-emerin (rabbit, abcam Ab40688, RRID: AB_2100059), anti-HA (mouse, Covance, MMS-101P, AB_2314672), anti-lamin A/C (mouse, ImmuQuest (IQ332 RRID 10660272)), anti-lamin B1 (rabbit, abcam ab16048, RRID: AB_443298), anti-lamin B2 (rabbit, abcam (ab151735) RRID: 2827514), anti-LAP1 (rabbit, abcam, ab86307 RRID: 2206124), anti-LBR (rabbit; Abnova, PAB15583; RRID: AB_10696691), mAB414 (mouse; Abcam, ab 24609; RRID: AB_448181), anti-H3 (rabbit, acam ab1791, RRID: 302613), anti-Tor1A (rabbit, Abexxa, abx001683), anti-Tor1B (rabbit, antibodies-online, ABIN1860834), anti-HSP60 (rabbit, abcam, ab45134, RRID 733033), anti-α-tubulin (mouse, Sigma, T5168, RRID 477579). Antibodies were raised in rabbits against purified, recombinant human LULL1. Antibodies against human SUN1 (Sosa et al., 2012), human SUN2 (Turgay et al., 2010) and anti-GFP (Turgay et al., 2014) have been previously described.

### Generation of LMN A/C knockout cells

*LMNA/C* KO cell lines were made using the CRISPR/Cas9 system. The guide RNA (gRNA) was designed using the CRISPR design web tools (http://www.e-crisp.org/E-CRISP/) and targeted exon 1 of the *LMNA/C* gene. The annealed guide RNA-encoding DNA oligos (5’-CACCGGGTGGCGCGCCGCTGGGACG-3’ and 5’-AAACCGTCCCAGCGGCGCGCCACCC-3’) were ligated into the pC2P vector (Welte et al., 2019), which encodes for hCas9 and possesses a puromycin resistance cassette. Cells were transfected with the resulting pC2P-gLMNA/C vector and selected with puromycin for 3 days. Surviving clones were expanded and screened for mutations in the *LMNA/C* gene by immunoblotting and PCR. PCR products covering the edited genomic region were sequenced and analyzed for indel mutations using the tide web tool (Brinkman et al., 2014) and manual inspection of the sequencing chromatograms.

### Cell culture and synchronization of cells

HeLa FRT/TetR, HeLa K, HCT116, and HepG2 were kind gifts from T. Mayer (Department of Biology, Konstanz), B. Vogelstein (Johns Hopkins, Baltimore), D. Gerlich (Institute of Molecular Biotechnology, Vienna), and S. Werner (Institute of Molecular Health Science, ETH Zurich). These cell lines were not further authenticated after obtaining them from the indicated sources. All cell lines were tested negative for mycoplasma using PCR-based testing. None of these cell lines was included in the list of commonly misidentified cell lines maintained by International Cell Line Authentication Committee. Cells were cultivated in DMEM containing 10% (v/v) FBS and 100 μg/ml penicillin/streptomycin at 37°C with 5% CO_2_ in a humidified incubator. Transient transfections were performed using X-tremeGene (Roche) and jetPRIME (PolyPlus) transfection reagent. Stable tetracycline-inducible HeLa cell lines were generated by integration of the respective constructs into HeLa FRT/TetR cells. Expression of the LAP1B-GFP, LAP1C-GFP, LAP2β-GFP, LAP1B(R563G)-GFP, LAP1B(E482A)-GFP, LAP1B(ΔCBR)-GFP, LAP1B(1-359)-GFP, LAP1B(S108E,T124E), Tor1B-GFP and Tor1B(E178Q)-GFP were induced with tetracycline as described in the figure legends. To synchronize HeLa cells in mitosis, a thymidine block was performed by adding 3 mM thymidine (Sigma-Aldrich) for 16 h, followed by release of the cells from G1/S arrest with fresh medium for 10 h. To arrest cells in prometaphase, 100 ng/ml nocodazole (Sigma-Aldrich) was added 8 h after thymidine release.

### Plasmid and siRNA transfections

siRNA transfections were performed using INTERFERin™ (Polyplus) as transfection reagent. Cells were treated with 20 nM siRNA for 48 h or 72 h. The following siRNAs were purchased:

si-CTR (AllStars negative control, Qiagen)

si-lamin A/C (CUGGACUUCCAGAAGAACA, Microsynth)

si-lamin B1 (UUCCGCCUCAGCCACUGGAAAU, Sigma)

si-lamin B2 (ACAACUCGGACAAGGAUC, Microsynth)

si-LAP1 CUCACUAAGUUUCCUGAGUUA, Microsynth)

si-LULL1 (CTGGTCCTGACTGTTCTGCTA, Microsynth)

si-Tor1A (CACCAAGTTAGATTATTACTA, Microsynth)

si-Tor1B (CTGTCGGAGTCTTCAATAATA, Microsynth)

### Immunofluorescence analysis

Cells grown on glass coverslips were washed with PBS and fixed for 10 min with either 4% PFA at RT or with methanol at −20°C. After PFA fixation, cells were permeabilized with 0.1% Triton X-100 in PBS for 10 min. Then, cells were blocked with 2% (w/v) BSA in PBS for at least 20 min, incubated with primary antibody for at least 1 h, washed three times with PBS and incubated with the respective secondary antibody for at least 45 min. Coverslips were washed again three times with PBS, incubated with 1 μg/ml Hoechst (Invitrogen) in PBS for 10 min and washed again. Coverslips were mounted on microscopic slides with Vectashield (Vector labs).

### Protein purification

Recombinant zz-His_6_, zz-LAP1B(98-136)-His_6_ wild-type and the respective 2E mutant (S108E, T124E) for EMSAs were produced in *E. coli* XL1(pBS161) (Kutay et al., 2000). Cells were lysed in 50 mM Tris pH 7.5, 700 mM NaCl, 2 mM MgCl_2_, 35 mM Imidazole, 2 mM beta-mercaptoethanol, 1 mg/ml lysozyme, 10 μg/ml DNase I, the recombinant proteins retrieved by immobilized nickel affinity chromatography, and eluted with 50 mM Tris pH 7.5, 350 mM NaCl, 2 mM MgCl_2_, 400 mM Imidazole, 2 mM beta-mercaptoethanol. The protein was rebuffered to 10 mM Tris pH 7.5, 200 mM NaCl, 1 mM EDTA, 1 mM DTT, 5% glycerol, 10% sucrose.

### Electrophoretic mobility shift assays (EMSAs) of DNA and mononucleosomes

DNA was produced as previously described (Hanson et al., 2004). In brief, a pUC19 plasmid carrying multiple copies of the Widom ‘601’ sequence, each flanked by EcoRV restriction sites, was purified from an 8 l culture of transformed *E. coli* DH5α cells (NEB). The ‘601’ fragments were released from the plasmid by digestion with EcoRV (NEB), isolated by incremental PEG precipitation, and further purified by ethanol-acetate precipitation and subsequent chloroform-phenol extraction. Histone octamers were prepared as described (Luger et al., 1999). We used wild-type histone proteins, except for H3.2 carrying a C110A mutation. Mononucleosomes were reconstituted through overnight logarithmic dialysis from high salt (10 mM Tris pH 7.5, 1 mM EDTA 2 M KCl) into low salt buffer (10 mM Tris pH 7.5, 1 mM EDTA, 10mM KCl), yielding a 22.6 nM mononucleosome solution based on DNA absorbance.

EMSAs were performed with 10 nM mononucleosomes or 10 nM ‘601’ DNA, preincubated with LAP1 protein derivatives at the indicated concentrations in a total volume of 10 μl for 10 min at 37°C (final buffer concentration 10 mM Tris pH 7.5, 1 mM EDTA, 150 mM KCl). Samples were put on ice, mixed with 5 μl ice-cold sample buffer (25% (w/v) sucrose, 0.01% (w/v) bromophenol blue) and loaded on a 5% polyacrylamide (29:1, BioRad) 0.5 x TBE gel, migrated for 1 h at 90 V in ice-cold 0.5 x TBE. DNA was stained with GelRed (Biotium) and imaged with a ChemiDoc MP(BioRad). Band intensities were quantified with ImageLab (BioRad). The intensities were normalized to the bands in the absence of added protein, and the averaged values of the replicates were plotted ± standard deviation. The data were evaluated with a Hill 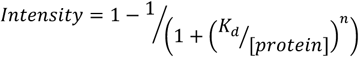 taking the standard deviation in account using IGOR (Wavemetrics).

### Image acquisition and live cell imaging

Confocal fluorescence images were acquired with LSM 880, LSM 780 microscope (ZEISS) or a Leica SP2 AOBS microscope. For all microscopes, 63× 1.4NA, oil plan-apochromat objectives were used. For time-lapse microscopy, cells were seeded into μ-slide 4-well ibidi chambers and incubated with 0.1 μM SiR-Hoechst 1 h prior to the experiment. Images were acquired at a Visitron spinning disk Nikon Eclipse T1 microscope, using a 60× 1.4 oil lens, in a 5% CO_2_, 37°C chamber. 15 z-stacks were acquired every 5 min for 8 to 12 h.

### FRAP

Cells were seeded into NuncTM Lab-TekTMII chambers, and transfected or induced one day before the experiment. For FRAP, either an LSM 710-FCS, LSM 780-FCS or LSM 880 Carl Zeiss microscope was used. Cells were kept in a humified chamber at 5% CO_2_ at 37°C. Time-lapse movies were recorded with a 63 × 1.4 NA oil DIC plan-apochromat immersion lens, and images were taken with a 4 × zoom. Three pre-bleach values were acquired, and then a 3.3 × 0.66 (2.2 μm^2^) rectangle at the NE was bleached (with 100% 488 nm laser intensity, and a range of 80-200 scanning iterations to achieve 90% bleaching). The fluorescence recovery was measured every 4 s for 57 cycles. The acquired FRAP data was normalized and fitted by a double exponential using the easyFRAP Matlab tool (Rapsomaniki et al, 2012).

### Statistical methods

Graphs were generated using Prism, which was further used to calculate the unpaired t-test and *p* values less than 0.001 are indicated by four asterisks (\*\*\*\**p* ≤ 0.0001), three asterisks (\*\*\**p* ≤ 0.001); *p* values less than 0.01 by two asterisks (\*\**p* ≤ 0.01); *p* values less than 0.05 by one asterisk (\**p* ≤ 0.05); and *p* values higher than 0.05 are marked as “not significant” (ns).

### Immunoblotting

Whole cell lysates were generated by resuspending cells in SDS sample buffer (75 mM Tris pH 6.8, 20% (v/v) glycerol, 4% (w/v) SDS, 50 mM DTT, 0.01% (w/v) bromophenole blue). Proteins were separated on 8-15% SDS gels and transferred to a nitrocellulose membrane (GE Healthcare) using the Trans-Blot SD Semi-Dry Transfer Cell (Bio-Rad). Membranes were incubated after blocking with 5% (w/v) dry milk in TBT with the primary antibody for at least 1 h. As secondary antibodies, rabbit anti-mouse or goat anti-rabbit horseradish-peroxidase-conjugated antibodies (Sigma) were used. Signals were captured by exposure to film or detected using a Fusion (Vilber) or an Odyssey (LI-COR) imaging system.

## Acknowledgements

We thank Dr. Sumit Pawar, Annamaria Gamper and Jasmin van den Heuvel for critical reading of the manuscript as well as the members of the Kutay lab for helpful discussions. Microscopy was performed on instruments of the ETHZ Microscopy Center Scope M. This work was funded by a grant of the Swiss National Science Foundation (SNSF) to U.K. (310030_184801), and by EPFL (B.F.).

---

**Figure 1, Supporting Figure 1.**
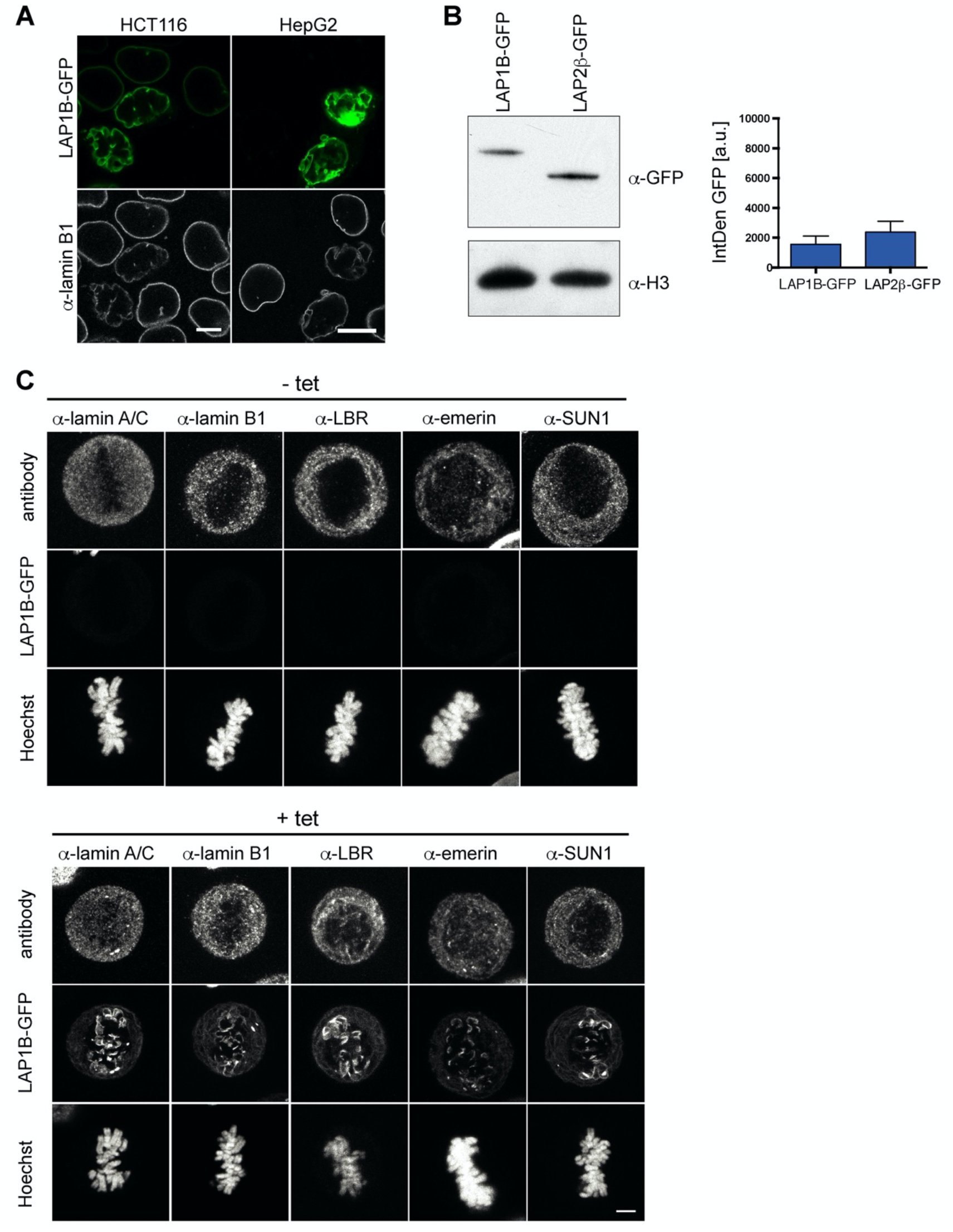
Interphase and mitotic localization of LAP1B-GFP and endogenous NE proteins. (A) Vector encoding LAP1B-GFP was transiently transfected into HCT116 or HepG2 cells. Cells were fixed after 48 h and analyzed by confocal microscopy. Scale bars, 10 μm. (B) Western blot analysis (left) and integrated GFP fluorescence intensities (right) in LAP1B-GFP and LAP2β-GFP-expressing cells of the experiment depicted in Fig. 1B. (C) Comparison of the metaphase localization of nuclear lamins and select INM proteins in an inducible LAP1B-GFP cell line without and with tetracycline (tet) induction. Maximum intensity projections (5 x 0.63 μm). Scale bar, 5 μm.

**Figure 1, Supporting Figure 2.**
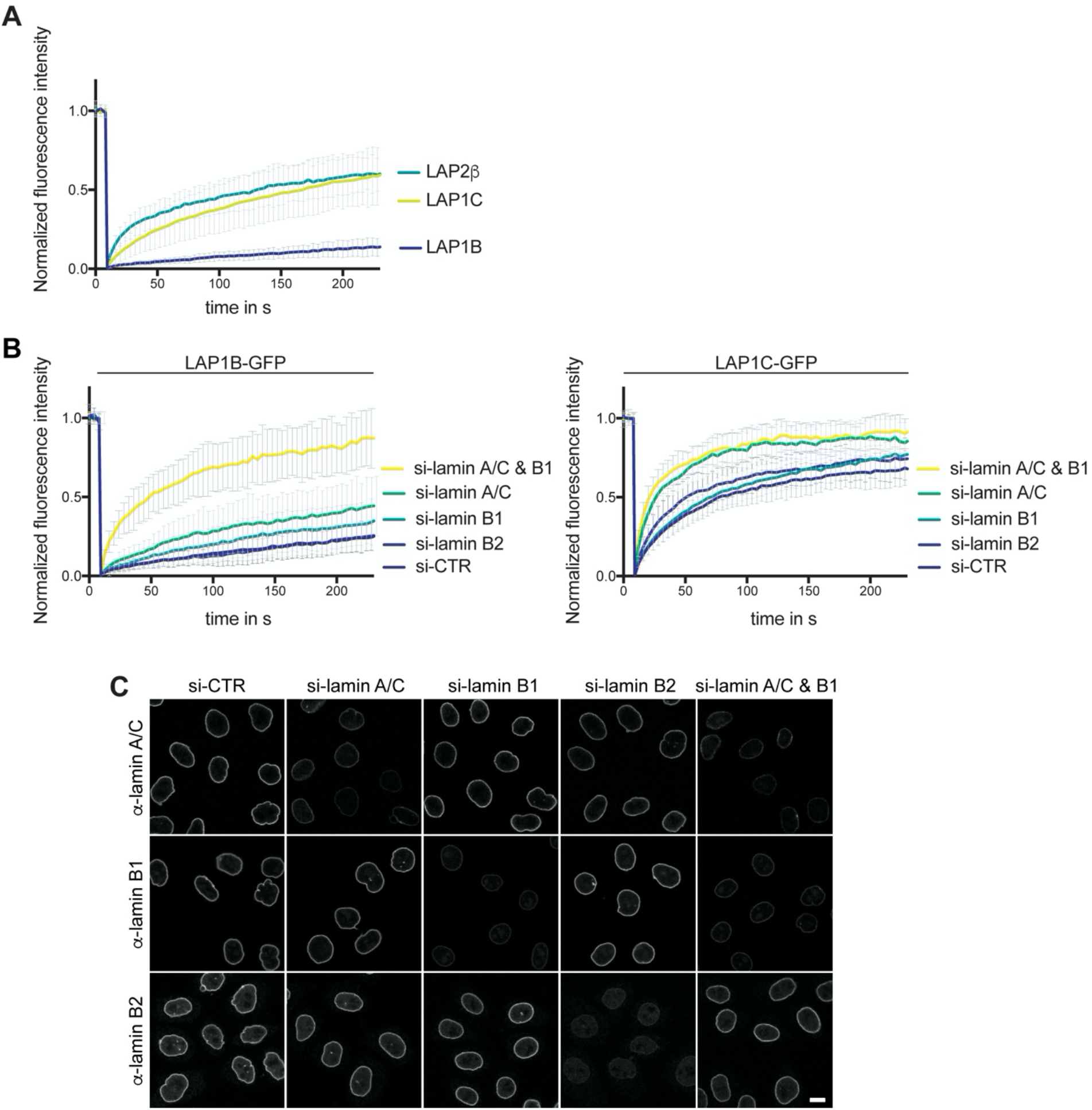
Lamin A/C and lamin B1 contribute to the retention of LAP1B at the NE. (A) FRAP analysis of LAP1B-GFP, LAP1C-GFP and LAP2β-GFP at the NE (N = 5, n > 18, mean +/- SD). (B) Diffusional mobility of LAP1B-GFP (N = 5, n > 26, mean +/- SD) and LAP1C-GFP (N = 4, n > 21, mean +/- SD) after RNAi-mediated depletion of lamins analyzed by FRAP (N = 5, n > 21, mean +/- SD). (C) Depletion of lamins by RNAi was efficient, as shown by immunofluorescence analysis of lamin A/C, lamin B1 and lamin B2. Scale bar, 10 μm.

**Figure 2, Supporting Figure 1.**
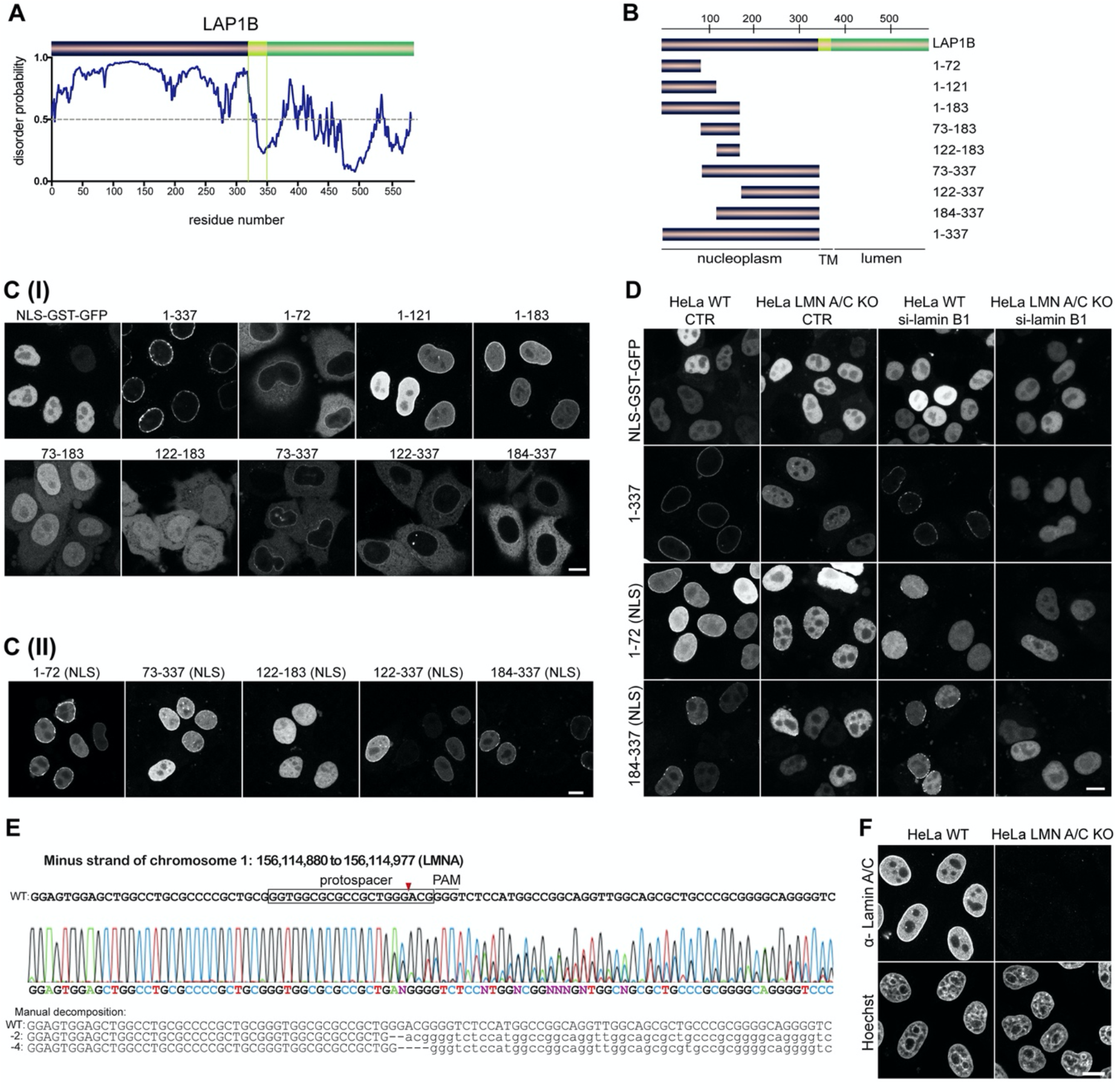
Delineation of the lamina-binding regions in LAP1B. (A) Scheme illustrating the disorder probability of LAP1. (B) LAP1B fragments used to delineate lamina-binding regions in the nucleoplasmic domain of LAP1B. (C) Localization of the indicated LAP1B derivatives fused to GST-GFP (I) or NLS-GST-GFP (II) in interphase HeLa cells. An NLS-GST-GFP fusion served as control. Note that enrichment of protein fragments at the nuclear rim reflects lamina association, as confirmed in panel D. Scale bars, 10 μm. (D) Localization of LAP1 fragments fused to GST-GFP-NLS in either wild-type or LMN A/C KO HeLa cells, with or without downregulation of lamin B1. Note that NE localization of LAP1(1-337) is strongly affected in LMN A/C KO cells, whereas the N-terminal fragment LAP1(1-72) is more strongly impaired in NE localization upon downregulation of lamin B1. Scale bar, 10 μm. (E) Characterization of LMN A/C knockout HeLa cells generated by CRISPR/Cas9 by DNA sequencing. (F) Analysis of LMN A/C knockout HeLa cells by immunofluorescence against lamin A/C. Scale bar, 10 μm.

**Figure 2, Supporting Figure 2.**
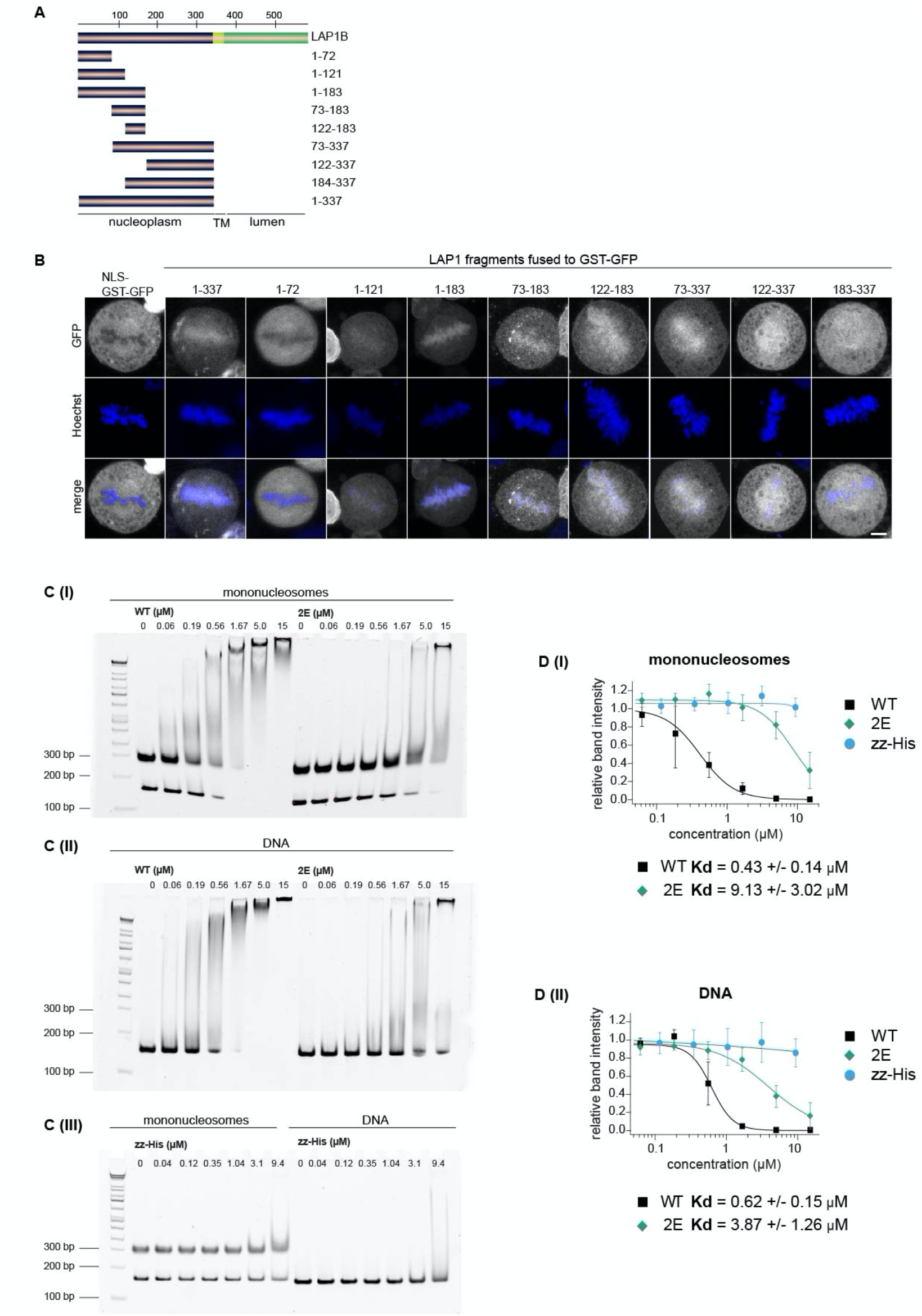
Delineation of the chromatin-binding region (CBR) of LAP1B. (A) LAP1B fragments used to map the minimal CBR in the nucleoplasmic domain of LAP1. (B) Localization of the indicated GST-GFP fusion proteins in synchronized HeLa cells during metaphase. DNA was stained with Hoechst. Maximum intensity projections (5 x 0.63 μm). Scale bar, 5 μm. (C) Electrophoretic mobility shift assays used for quantitative analysis of DNA and nucleosome binding of the isolated recombinant CBR of LAP1B. Increasing concentrations of zz-LAP1(98-136), zz-LAP1(98-136; S108E, T124E (‘2E’)) (I and II) or zz-His (III) were incubated with either ‘601’ DNA or reconstituted mononucleosomes and subjected to native gel electrophoresis. DNA was stained with GelRed. (D) Quantification of experiments in C based on integration of gel-band intensities from 3 independent experiments (N = 3), (I) for binding to mononucleosomes, (II) to DNA. Normalized band intensities (mean +/- SD) were plotted over the protein concentration and K_d_ values calculated using the Hill equation.

**Figure 2, Supporting Figure 3.**
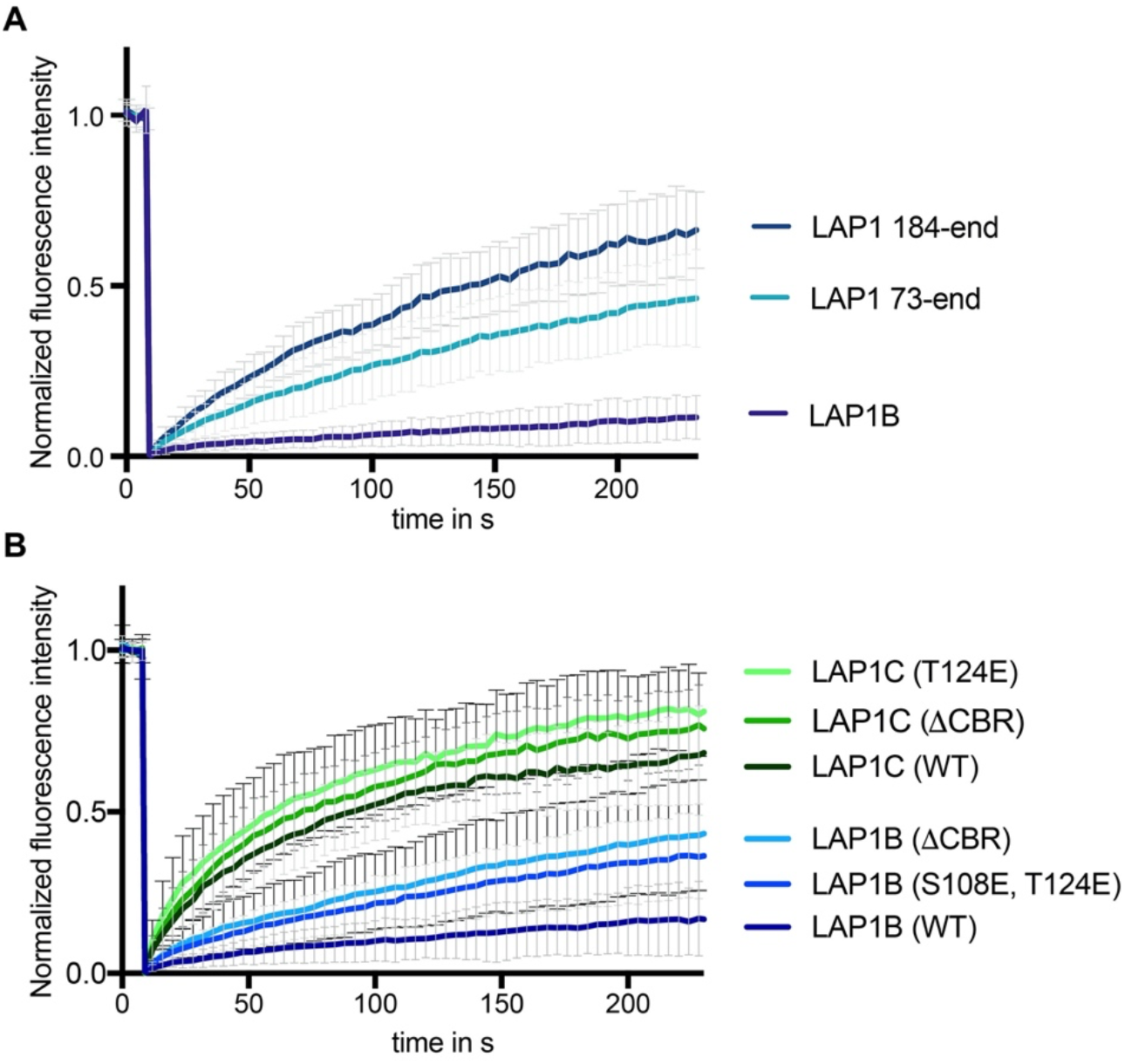
The CBR of LAP1 contributes to its retention at the INM. (A) FRAP analysis of LAP1B-GFP wild-type in comparison to truncation constructs lacking the N-terminal lamin-binding domain (LAP1(73-end)) or an extended part including the CBR (LAP1(184-end)); (N = 3, n > 20, mean +/- SD). Scale bar, 5 µm. (B) FRAP analysis of LAP1B and LAP1C-GFP and the indicated mutants of the chromatin-binding region (N=3, n > 14, mean +/- SD).

**Figure 3, Supporting Figure 1.**
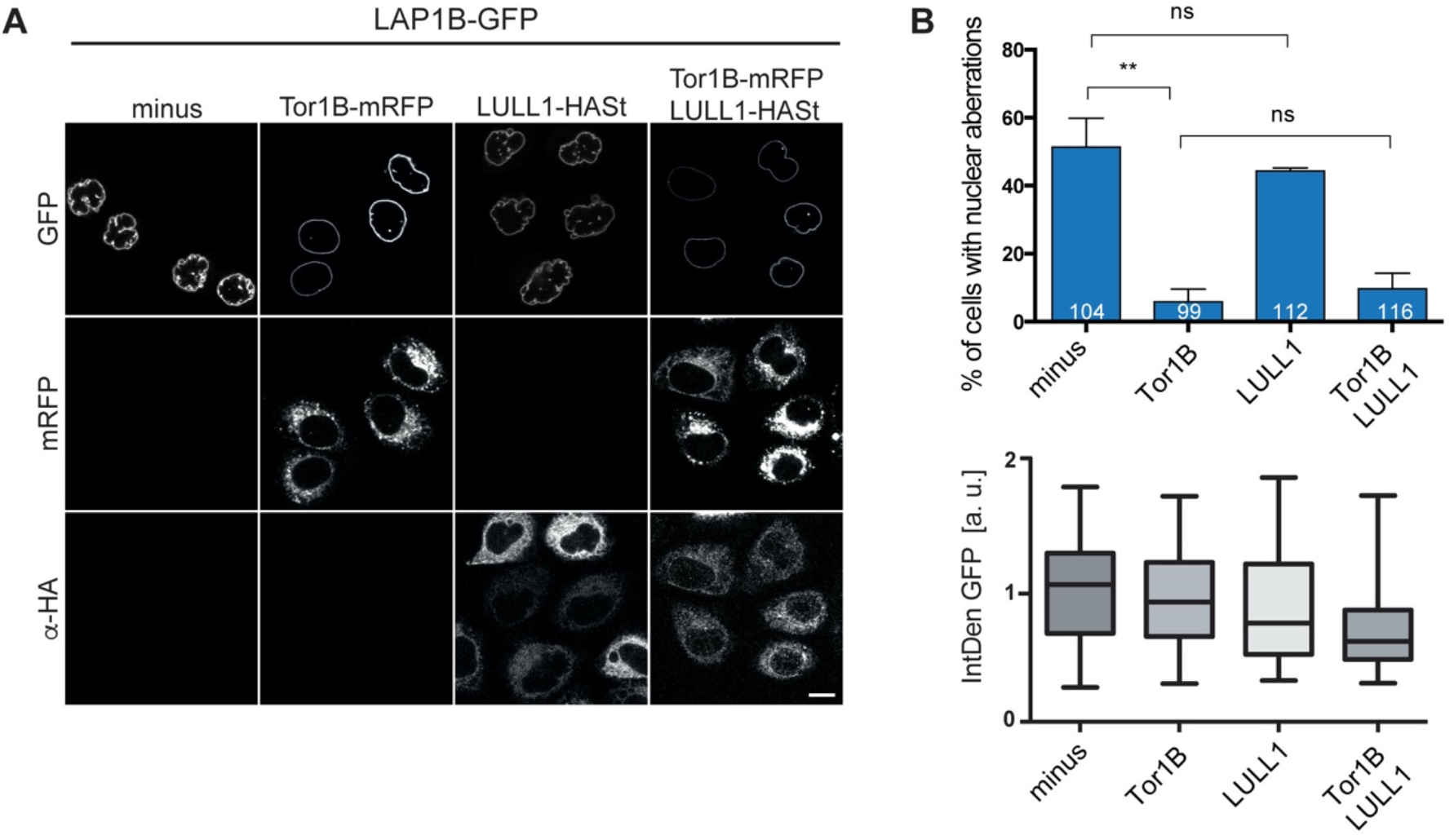
NE aberrations induced by LAP1B cannot be rescued by LULL1 co-expression. (A) HeLa cells were transfected with constructs encoding for LAP1B-GFP, either alone or together with LULL1-HASt and Tor1B-mRFP expression vectors. Cell were fixed 48 h after transfection and nuclear morphology analyzed by confocal microscopy. Scale bar, 10 μm. (B) Quantification of NE aberrations in cells with similar LAP1B-GFP expression levels (N = 3, n > 99, mean +/- SEM).

**Figure 6, Supporting Figure 1.**
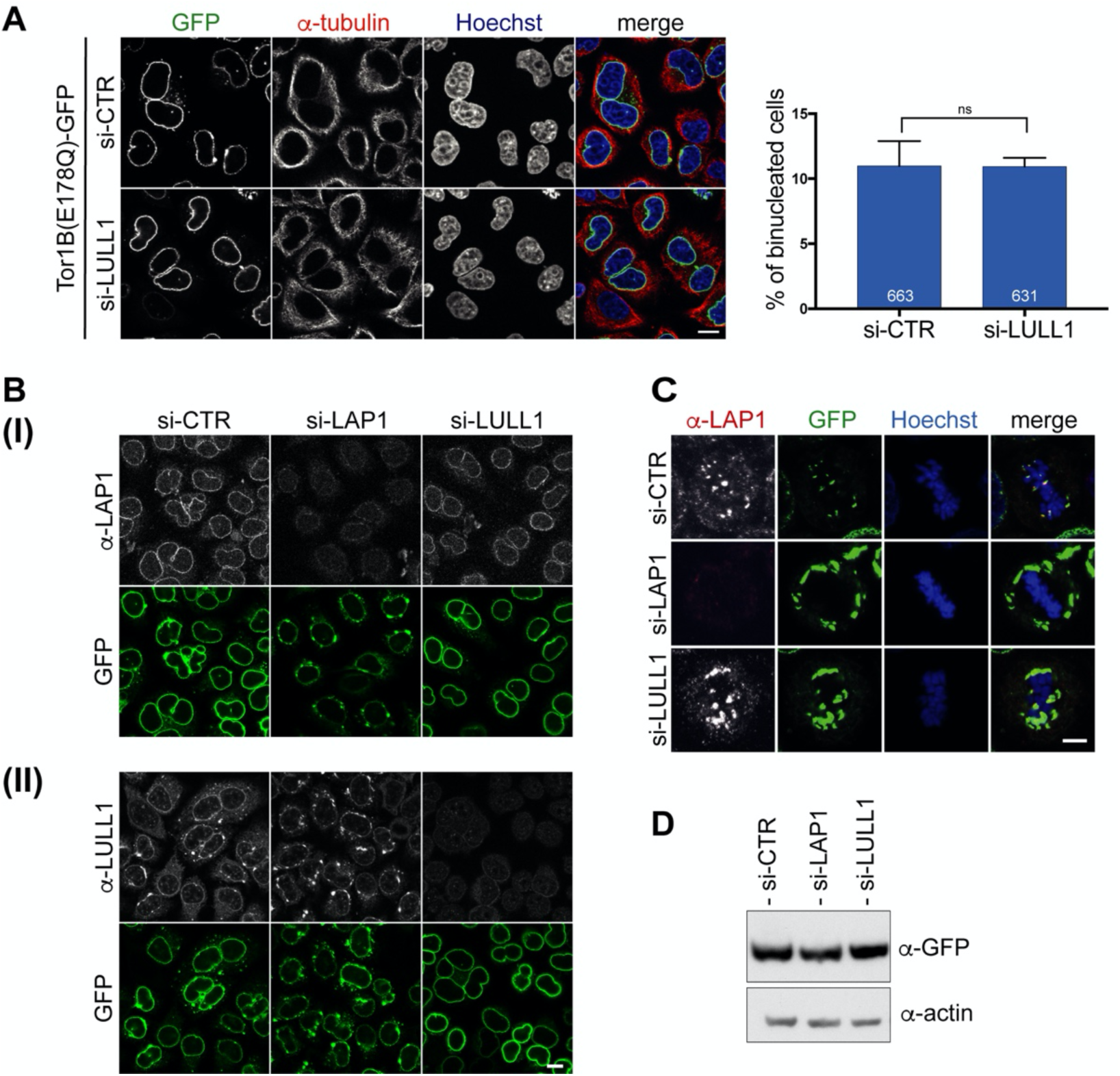
Downregulation of LULL1 does not prevent Tor1B(E178Q)-induced binucleation. (A) Expression of Tor1B(E178Q)-GFP was induced for 48 h in control or LULL1-depleted cells. Binucleation was quantified as in Figs. 5-7 (N = 3, n > 631, mean+/- SEM). Scale bar, 10 μm. (A) Depletion of LAP1 (I) and LULL1 (II) was controlled by immunofluorescence. Tor1B(E178Q)-GFP was expressed at comparable levels as shown by immunoblotting in panel D. (B) Localization of endogenous LAP1 at metaphase chromatin upon overexpression of Tor1B(E178Q)-GFP disappears upon LAP1 depletion, but is not influenced by LULL1 depletion. Representative confocal images (maximal z-projections: 5 x 0.63 μm) of metaphase HeLa cells with TorsinB-E178Q-GFP expression. Localization of LAP1 was analyzed by immunofluorescence staining. Scale bar, 5 μm. (C) Western blot analysis of Tor1B(E178Q)-GFP levels after LAP1 or LULL1 depletion.

**Figure 6, Supporting Figure 2.**
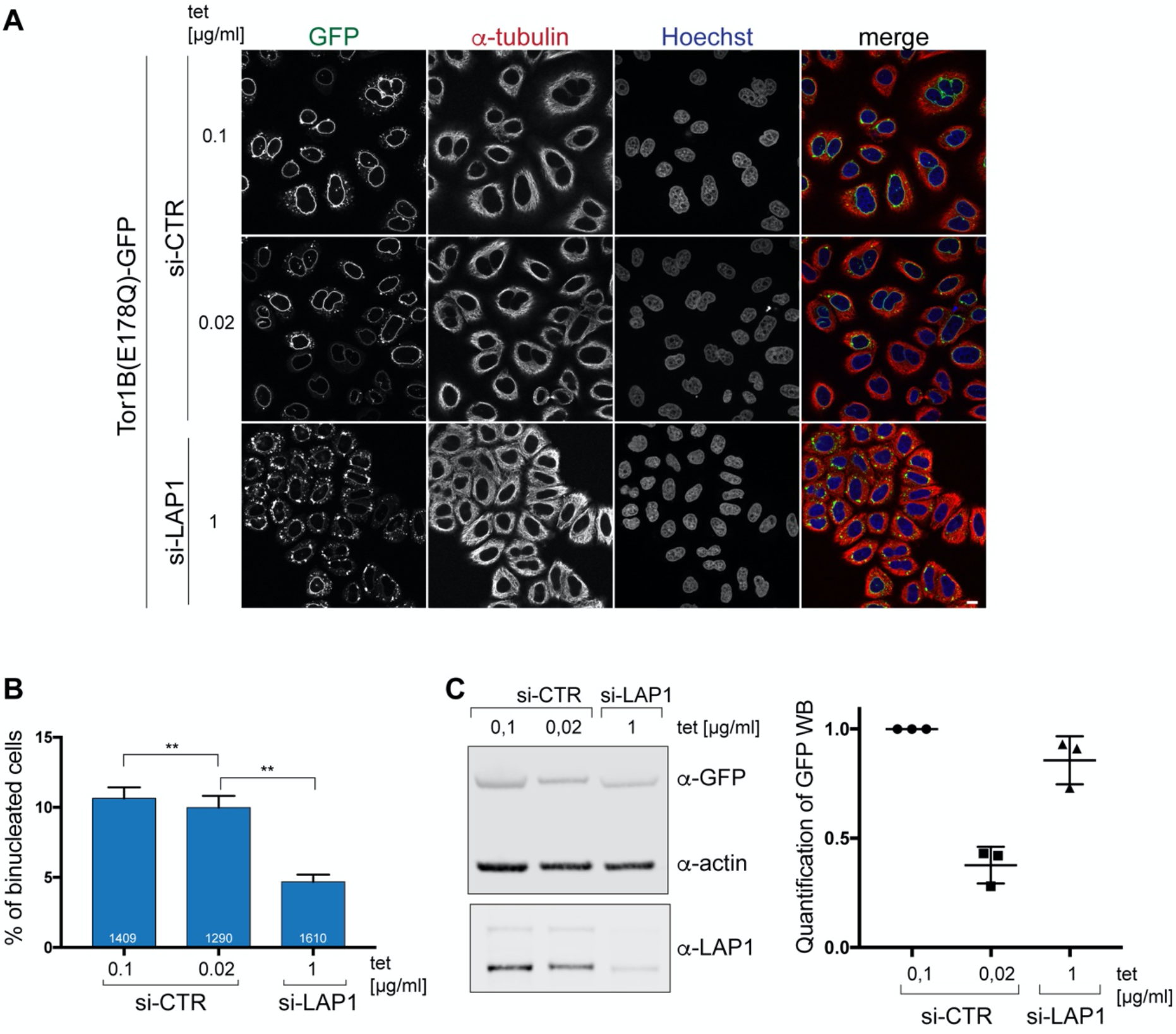
Expression of a dominant-negative Torsin1B variant leads to an increase in binucleated cells in a LAP1-dependent manner. (A) Expression of Tor1B(E178Q)-GFP was induced for 48 h in LAP1-depleted cells (1 μg/ml tet) and control siRNA-treated cells (0.1 μg/ml and 0.02 μg/ml tet). Scale bar, 10 μm. (B) Quantification of binucleated cells (N = 3, n > 631, mean+/- SEM). (C) Quantitative Western blot analysis of Tor1B(E178Q)-GFP levels relative to actin for depicted conditions.

## References

Barton LJ, Soshnev AA, Geyer PK (2015) Networking in the nucleus: a spotlight on LEM-domain proteins. Curr Opin Cell Biol 34: 1–8

Boni A, Politi AZ, Strnad P, Xiang W, Hossain MJ, Ellenberg J (2015) Live imaging and modeling of inner nuclear membrane targeting reveals its molecular requirements in mammalian cells. J Cell Biol 209: 705–20

Brachner A, Foisner R (2011) Evolvement of LEM proteins as chromatin tethers at the nuclear periphery. Biochem Soc Trans 39: 1735–41

Brinkman EK, Chen T, Amendola M, van Steensel B (2014) Easy quantitative assessment of genome editing by sequence trace decomposition. Nucleic Acids Res 42: e168

Brown RS, Zhao C, Chase AR, Wang J, Schlieker C (2014) The mechanism of Torsin ATPase activation. Proc Natl Acad Sci U S A 111: E4822–31

Cancer Genome Atlas N (2012) Comprehensive molecular characterization of human colon and rectal cancer. Nature 487: 330–7

Cascalho A, Jacquemyn J, Goodchild RE (2017) Membrane defects and genetic redundancy: Are we at a turning point for DYT1 dystonia? Mov Disord 32: 371–381

Champion L, Linder MI, Kutay U (2017) Cellular Reorganization during Mitotic Entry. Trends Cell Biol 27: 26–41

Champion L, Pawar S, Luithle N, Ungricht R, Kutay U (2019) Dissociation of membrane-chromatin contacts is required for proper chromosome segregation in mitosis. Mol Biol Cell 30: 427–440

Chase AR, Laudermilch E, Wang J, Shigematsu H, Yokoyama T, Schlieker C (2017) Dynamic functional assembly of the Torsin AAA+ ATPase and its modulation by LAP1. Mol Biol Cell 28: 2765–2772

Chen P, Burdette AJ, Porter JC, Ricketts JC, Fox SA, Nery FC, Hewett JW, Berkowitz LA, Breakefield XO, Caldwell KA, Caldwell GA (2010) The early-onset torsion dystonia-associated protein, torsinA, is a homeostatic regulator of endoplasmic reticulum stress response. Hum Mol Genet 19: 3502–15

Chow KH, Factor RE, Ullman KS (2012) The nuclear envelope environment and its cancer connections. Nat Rev Cancer 12: 196–209

Demircioglu FE, Sosa BA, Ingram J, Ploegh HL, Schwartz TU (2016) Structures of TorsinA and its disease-mutant complexed with an activator reveal the molecular basis for primary dystonia. Elife 5

Dephoure N, Zhou C, Villen J, Beausoleil SA, Bakalarski CE, Elledge SJ, Gygi SP (2008) A quantitative atlas of mitotic phosphorylation. Proc Natl Acad Sci U S A 105: 10762–7

Dorboz I, Coutelier M, Bertrand AT, Caberg JH, Elmaleh-Berges M, Laine J, Stevanin G, Bonne G, Boespflug-Tanguy O, Servais L (2014) Severe dystonia, cerebellar atrophy, and cardiomyopathy likely caused by a missense mutation in TOR1AIP1. Orphanet J Rare Dis 9: 174

Ellenberg J, Siggia ED, Moreira JE, Smith CL, Presley JF, Worman HJ, Lippincott-Schwartz J (1997) Nuclear membrane dynamics and reassembly in living cells: targeting of an inner nuclear membrane protein in interphase and mitosis. J Cell Biol 138: 1193–206

Erkmann JA, Wagner EJ, Dong J, Zhang Y, Kutay U, Marzluff WF (2005) Nuclear import of the stem-loop binding protein and localization during the cell cycle. Mol Biol Cell 16: 2960–71

Fichtman B, Zagairy F, Biran N, Barsheshet Y, Chervinsky E, Ben Neriah Z, Shaag A, Assa M, Elpeleg O, Harel A, Spiegel R (2019) Combined loss of LAP1B and LAP1C results in an early onset multisystemic nuclear envelopathy. Nat Commun 10: 605

Foisner R, Gerace L (1993) Integral membrane proteins of the nuclear envelope interact with lamins and chromosomes, and binding is modulated by mitotic phosphorylation. Cell 73: 1267–79

Goodchild RE, Buchwalter AL, Naismith TV, Holbrook K, Billion K, Dauer WT, Liang CC, Dear ML, Hanson PI (2015) Access of torsinA to the inner nuclear membrane is activity dependent and regulated in the endoplasmic reticulum. J Cell Sci 128: 2854–65

Goodchild RE, Dauer WT (2005) The AAA+ protein torsinA interacts with a conserved domain present in LAP1 and a novel ER protein. J Cell Biol 168: 855–62

Goodchild RE, Kim CE, Dauer WT (2005) Loss of the dystonia-associated protein torsinA selectively disrupts the neuronal nuclear envelope. Neuron 48: 923–32

Grillet M, Dominguez Gonzalez B, Sicart A, Pottler M, Cascalho A, Billion K, Hernandez Diaz S, Swerts J, Naismith TV, Gounko NV, Verstreken P, Hanson PI, Goodchild RE (2016) Torsins Are Essential Regulators of Cellular Lipid Metabolism. Dev Cell 38: 235–47

Guelen L, Pagie L, Brasset E, Meuleman W, Faza MB, Talhout W, Eussen BH, de Klein A, Wessels L, de Laat W, van Steensel B (2008) Domain organization of human chromosomes revealed by mapping of nuclear lamina interactions. Nature 453: 948–51

Guttinger S, Laurell E, Kutay U (2009) Orchestrating nuclear envelope disassembly and reassembly during mitosis. Nat Rev Mol Cell Biol 10: 178–91

Hanson BL, Alexander C, Harp JM, Bunick GJ (2004) Preparation and crystallization of nucleosome core particle. Methods Enzymol 375: 44–62

Hanson PI, Whiteheart SW (2005) AAA+ proteins: have engine, will work. Nat Rev Mol Cell Biol 6: 519–29

Hirano Y, Hizume K, Kimura H, Takeyasu K, Haraguchi T, Hiraoka Y (2012) Lamin B receptor recognizes specific modifications of histone H4 in heterochromatin formation. J Biol Chem 287: 42654–63

Holmer L, Worman HJ (2001) Inner nuclear membrane proteins: functions and targeting. Cell Mol Life Sci 58: 1741–7

Itzhak DN, Tyanova S, Cox J, Borner GH (2016) Global, quantitative and dynamic mapping of protein subcellular localization. Elife 5

Jokhi V, Ashley J, Nunnari J, Noma A, Ito N, Wakabayashi-Ito N, Moore MJ, Budnik V (2013) Torsin Mediates Primary Envelopment of Large Ribonucleoprotein Granules at the Nuclear Envelope. In Cell reports,

Jungwirth M, Dear ML, Brown P, Holbrook K, Goodchild R (2010) Relative tissue expression of homologous torsinB correlates with the neuronal specific importance of DYT1 dystonia-associated torsinA. Hum Mol Genet 19: 888–900

Kayman-Kurekci G, Talim B, Korkusuz P, Sayar N, Sarioglu T, Oncel I, Sharafi P, Gundesli H, Balci-Hayta B, Purali N, Serdaroglu-Oflazer P, Topaloglu H, Dincer P (2014) Mutation in TOR1AIP1 encoding LAP1B in a form of muscular dystrophy: a novel gene related to nuclear envelopathies. Neuromuscul Disord 24: 624–33

Kim CE, Perez A, Perkins G, Ellisman MH, Dauer WT (2010) A molecular mechanism underlying the neural-specific defect in torsinA mutant mice. Proc Natl Acad Sci U S A 107: 9861–6

Krauthammer M, Kong Y, Ha BH, Evans P, Bacchiocchi A, McCusker JP, Cheng E, Davis MJ, Goh G, Choi M, Ariyan S, Narayan D, Dutton-Regester K, Capatana A, Holman EC, Bosenberg M, Sznol M, Kluger HM, Brash DE, Stern DF et al. (2012) Exome sequencing identifies recurrent somatic RAC1 mutations in melanoma. Nat Genet 44: 1006–14

Kutay U, Hartmann E, Treichel N, Calado A, Carmo-Fonseca M, Prehn S, Kraft R, Gorlich D, Bischoff FR (2000) Identification of two novel RanGTP-binding proteins belonging to the importin beta superfamily. J Biol Chem 275: 40163–8

Kutay U, Izaurralde E, Bischoff FR, Mattaj IW, Gorlich D (1997) Dominant-negative mutants of importin-beta block multiple pathways of import and export through the nuclear pore complex. EMBO J 16: 1153–63

Laudermilch E, Schlieker C (2016) Torsin ATPases: structural insights and functional perspectives. Curr Opin Cell Biol 40: 1–7

Levine MS, Holland AJ (2018) The impact of mitotic errors on cell proliferation and tumorigenesis. Genes Dev 32: 620–638

Luger K, Rechsteiner TJ, Richmond TJ (1999) Preparation of nucleosome core particle from recombinant histones. Methods Enzymol 304: 3–19

Maric M, Shao J, Ryan RJ, Wong C-S, Gonzalez-Alegre P, Roller RJ (2011) A functional role for TorsinA in herpes simplex virus 1 nuclear egress. In J Virol, pp 9667-9679. American Society for Microbiology

Nery FC, Armata IA, Farley JE, Cho JA, Yaqub U, Chen P, da Hora CC, Wang Q, Tagaya M, Klein C, Tannous B, Caldwell KA, Caldwell GA, Lencer WI, Ye Y, Breakefield XO (2011) TorsinA participates in endoplasmic reticulum-associated degradation. Nat Commun 2: 393

Olivares AO, Baker TA, Sauer RT (2016) Mechanistic insights into bacterial AAA+ proteases and protein-remodelling machines. Nat Rev Microbiol 14: 33–44

Ozelius LJ, Hewett JW, Page CE, Bressman SB, Kramer PL, Shalish C, de Leon D, Brin MF, Raymond D, Corey DP, Fahn S, Risch NJ, Buckler AJ, Gusella JF, Breakefield XO (1997) The early-onset torsion dystonia gene (DYT1) encodes an ATP-binding protein. In Nat Genet, pp 40–48.

Pappas SS, Liang CC, Kim S, Rivera CO, Dauer WT (2018) TorsinA dysfunction causes persistent neuronal nuclear pore defects. Hum Mol Genet 27: 407–420

Powell L, Burke B (1990) Internuclear exchange of an inner nuclear membrane protein (p55) in heterokaryons: in vivo evidence for the interaction of p55 with the nuclear lamina. J Cell Biol 111: 2225–34

Rampello AJ, Laudermilch E, Vishnoi N, Prohet SM, Shao L, Zhao C, Lusk CP, Schlieker C (2019) Torsin ATPases are required to complete nuclear pore complex biogenesis in interphase. bioRxiv

Rebelo S, da Cruz ESEF, da Cruz ESOA (2015) Genetic mutations strengthen functional association of LAP1 with DYT1 dystonia and muscular dystrophy. Mutat Res Rev Mutat Res 766: 42–7

Rose AE, Brown RS, Schlieker C (2015) Torsins: not your typical AAA+ ATPases. Crit Rev Biochem Mol Biol 50: 532–49

Rose AE, Zhao C, Turner EM, Steyer AM, Schlieker C (2014) Arresting a Torsin ATPase reshapes the endoplasmic reticulum. J Biol Chem 289: 552–64

Samwer M, Schneider MWG, Hoefler R, Schmalhorst PS, Jude JG, Zuber J, Gerlich DW (2017) DNA Cross-Bridging Shapes a Single Nucleus from a Set of Mitotic Chromosomes. Cell 170: 956–972 e23

Santos M, Costa P, Martins F, da Cruz e Silva EF, da Cruz e Silva OA, Rebelo S (2015) LAP1 is a crucial protein for the maintenance of the nuclear envelope structure and cell cycle progression. Mol Cell Biochem 399: 143–53

Santos M, Domingues SC, Costa P, Muller T, Galozzi S, Marcus K, da Cruz e Silva EF, da Cruz e Silva OA, Rebelo S (2014) Identification of a novel human LAP1 isoform that is regulated by protein phosphorylation. PLoS One 9: e113732

Santos M, Rebelo S, Van Kleeff PJ, Kim CE, Dauer WT, Fardilha M, da Cruz ESOA, da Cruz ESEF (2013) The nuclear envelope protein, LAP1B, is a novel protein phosphatase 1 substrate. PLoS One 8: e76788

Senior A, Gerace L (1988) Integral membrane proteins specific to the inner nuclear membrane and associated with the nuclear lamina. J Cell Biol 107: 2029–36

Shin JY, Dauer WT, Worman HJ (2014) Lamina-associated polypeptide 1: protein interactions and tissue-selective functions. Semin Cell Dev Biol 29: 164–8

Shin JY, Hernandez-Ono A, Fedotova T, Ostlund C, Lee MJ, Gibeley SB, Liang CC, Dauer WT, Ginsberg HN, Worman HJ (2019) Nuclear envelope-localized torsinA-LAP1 complex regulates hepatic VLDL secretion and steatosis. J Clin Invest 130: 4885–4900

Snider J, Thibault G, Houry WA (2008) The AAA+ superfamily of functionally diverse proteins. Genome Biol 9: 216

Solovei I, Wang AS, Thanisch K, Schmidt CS, Krebs S, Zwerger M, Cohen TV, Devys D, Foisner R, Peichl L, Herrmann H, Blum H, Engelkamp D, Stewart CL, Leonhardt H, Joffe B (2013) LBR and lamin A/C sequentially tether peripheral heterochromatin and inversely regulate differentiation. Cell 152: 584–98

Sosa BA, Demircioglu FE, Chen JZ, Ingram J, Ploegh HL, Schwartz TU (2014) How lamina-associated polypeptide 1 (LAP1) activates Torsin. Elife 3: e03239

Sosa BA, Rothballer A, Kutay U, Schwartz TU (2012) LINC complexes form by binding of three KASH peptides to domain interfaces of trimeric SUN proteins. Cell 149: 1035–47

Tanaka K, Goto H, Nishimura Y, Kasahara K, Mizoguchi A, Inagaki M (2018) Tetraploidy in cancer and its possible link to aging. Cancer Sci 109: 2632–2640

Titos I, Ivanova T, Mendoza M (2014) Chromosome length and perinuclear attachment constrain resolution of DNA intertwines. J Cell Biol 206: 719–33

Turgay Y, Champion L, Balazs C, Held M, Toso A, Gerlich DW, Meraldi P, Kutay U (2014) SUN proteins facilitate the removal of membranes from chromatin during nuclear envelope breakdown. J Cell Biol 204: 1099–109

Turgay Y, Ungricht R, Rothballer A, Kiss A, Csucs G, Horvath P, Kutay U (2010) A classical NLS and the SUN domain contribute to the targeting of SUN2 to the inner nuclear membrane. EMBO J 29: 2262–75

Turner EM, Brown RS, Laudermilch E, Tsai PL, Schlieker C (2015) The Torsin Activator LULL1 Is Required for Efficient Growth of Herpes Simplex Virus 1. J Virol 89: 8444–52

Ulbert S, Platani M, Boue S, Mattaj IW (2006) Direct membrane protein-DNA interactions required early in nuclear envelope assembly. J Cell Biol 173: 469–76

Ungricht R, Klann M, Horvath P, Kutay U (2015) Diffusion and retention are major determinants of protein targeting to the inner nuclear membrane. J Cell Biol 209: 687–703

Ungricht R, Kutay U (2015) Establishment of NE asymmetry-targeting of membrane proteins to the inner nuclear membrane. Curr Opin Cell Biol 34: 135–41

Ungricht R, Kutay U (2017) Mechanisms and functions of nuclear envelope remodelling. Nat Rev Mol Cell Biol 18: 229–245

Vander Heyden AB, Naismith TV, Snapp EL, Hodzic D, Hanson PI (2009) LULL1 retargets TorsinA to the nuclear envelope revealing an activity that is impaired by the DYT1 dystonia mutation. Mol Biol Cell 20: 2661–72

Weibezahn J, Schlieker C, Bukau B, Mogk A (2003) Characterization of a trap mutant of the AAA+ chaperone ClpB. J Biol Chem 278: 32608–17

Welte T, Tuck AC, Papasaikas P, Carl SH, Flemr M, Knuckles P, Rankova A, Buhler M, Grosshans H (2019) The RNA hairpin binder TRIM71 modulates alternative splicing by repressing MBNL1. Genes Dev 33: 1221–1235

Yang L, Guan T, Gerace L (1997) Integral membrane proteins of the nuclear envelope are dispersed throughout the endoplasmic reticulum during mitosis. J Cell Biol 137: 1199–210

Zhao C, Brown RS, Chase AR, Eisele MR, Schlieker C (2013) Regulation of Torsin ATPases by LAP1 and LULL1. Proc Natl Acad Sci U S A 110: E1545–54

Zuleger N, Kelly DA, Richardson AC, Kerr AR, Goldberg MW, Goryachev AB, Schirmer EC (2011) System analysis shows distinct mechanisms and common principles of nuclear envelope protein dynamics. J Cell Biol 193: 109–23

